# Syntenic lncRNAs exhibit DNA regulatory functions with sequence evolution

**DOI:** 10.1101/2024.04.26.588027

**Authors:** Gyan Ranjan, Vinod Scaria, Sridhar Sivasubbu

## Abstract

Syntenic long non-coding RNAs (lncRNAs) often show limited sequence conservation across species, prompting concern in the field. This study delves into functional signatures of syntenic lncRNAs between humans and zebrafish. Syntenic lncRNAs have high expression in zebrafish and ∼90% near protein-coding genes in sense or antisense orientation. During early zebrafish development and human embryonic stem cells (H1-hESC), are enriched with cis-regulatory repressor signatures, influencing development-associated genes. In later zebrafish developmental stages and specific human cell lines, these lncRNAs serve as enhancers or transcription-start-sites(TSS) for protein-coding. Analysis of Transposable Elements (TEs) in syntenic lncRNA sequence divergence unveils intriguing patterns, human lncRNAs show enrichment in simple repeat elements, while zebrafish counterparts exhibit LTR element enrichment. This sequence evolution, possibly stemming from post-rearrangement mutations, enhances DNA elements or cis-regulatory functions. It may also contribute to vertebrate innovation by creating novel TF binding sites within the locus. This study sheds light on the conserved functionality of syntenic lncRNAs through DNA elements, emphasizing their role across species despite sequence divergence.

## Introduction

Long non-coding RNAs (lncRNAs) are important regulatory molecules with relatively unknown evolutionary significance across species. Researchers explore their functions and conservation using four key approaches: sequence motif, structural, syntenic, and functional conservation. (Diederichs, 2014; Ranjan et al, 2021; Szcześniak et al, 2021; Mattick et al, 2023). Over the past decade, several investigations have utilized a syntenic-based conservation analysis strategy, which is per se defined as maintaining the relative gene order and arrangement across various species throughout evolution. This approach has been employed to pinpoint the conserved lncRNAs among distinct organisms, offering perspectives into their potential functional importance and evolutionary persistence (Liu et al, 2018; Ranjan et al, 2021; Zhou et al, 2023). Recently, zebrafish emerged as a powerful model for reverse genetic studies of lncRNAs with unknown functions (Chen et al, 2018). This is due to its advantageous characteristics, including high fecundity, external fertilization with optical transparency, low-cost maintenance, and a comprehensively sequenced genome bearing significant similarity to the human genome (Howe et al, 2013; Patowary et al, 2013). The roles of syntenic lncRNA in early developmental processes have been substantiated through several candidate investigations, including studies involving *Megamind, tie1-AS, Alien, Terminator, Punisher, sox2-ot*, and more in zebrafish (Li et al, 2010; Ulitsky et al, 2011; Kurian et al, 2015; Ranjan et al, 2021). These findings underscore its significance as a valuable model system to functionally understand the role of this syntenic conserved lncRNA in complex systems. Although a large proportion of these lncRNA remains uncharacterized, their involvement in a complex system such as the development of zebrafish or humans remains unclear (Mattick et al, 2023). Leveraging extensive multi-omics datasets, we possess the capability to thoroughly analyze and comprehend the functional attributes of syntenic lncRNA, shedding light on their potential roles and contributions.

The lncRNAs have been noted to exert regulatory influence on their targets either in a *cis* or *trans* manner. Furthermore, these regulatory mechanisms employed by lncRNAs predominantly fall into three distinct functional modalities inherent within the lncRNAs-the RNA molecule, the regulatory elements within the lncRNA loci, or the act of transcription (Kopp & Mendell, 2018). In recent times, numerous investigations have emphasized the propensity of *cis*-regulating lncRNAs to manifest a distinctive DNA regulatory signature within the gene loci, playing pivotal roles in modulating the expression of neighboring protein-coding genes (Latos et al, 2012; Alexanian et al, 2017; Cho et al, 2018; Furlan et al, 2018; Ritter et al, 2019; Rom et al, 2019; Ali & Grote, 2020; Gil et al, 2023). These regulatory signatures encompass a spectrum of functions, including chromatin remodeling, enhancer activation, repressor activity, and epigenetic modulation. A limited number of lncRNAs have demonstrated the capacity for multi-modal functionality within their loci. One such instance is the *Haunt* lncRNA, where distinct functions are attributed to the mature RNA and DNA elements existing within the lncRNA locus (Yin et al, 2015). Another illustration is the *AIRN* lncRNA, wherein transcription itself regulates local gene expression, while the mature RNA participates in recruiting chromatin modifiers to induce silencing of neighboring genes (Latos et al, 2012; Santoro et al, 2013; Andergassen et al, 2019). Recently studied lncRNA *Tug1* locus suggested that it contains two distinct non-coding functions, acting as a *cis* DNA repressor and a *trans*-acting lncRNA, with a potential third protein-coding role (Lewandowski et al, 2020). Despite an extensive array of uncharacterized lncRNAs, the regulatory mechanisms underlying this specific lncRNA class remain enigmatic.

An intriguing observation revealed that a significant majority of human syntenic conserved lncRNAs indeed exhibited sequence conservation across the mammalian species but showed no sequence conservation in zebrafish and stickleback (Kapusta & Feschotte, 2014; Hezroni et al, 2015; Toiber et al, 2017). Moreover, low or no sequence conservation has been a characteristic feature of lncRNAs and multiple candidates have demonstrated that syntenic-conserved lncRNAs do not exhibit sequence similarity across species (Hezroni et al, 2015; Kurian et al, 2015; Alexanian et al, 2017; Ranjan et al, 2021; Mattick et al, 2023; Zhou et al, 2023). Multiple evolutionary models have been previously investigated suggesting these sequence discrepancies between humans and zebrafish may be attributed to a range of genomic rearrangement events. These events encompass genomic translocations, transpositions, inversions, and rearrangements, potentially leading to gene duplications and transposon-mediated alterations, acting as genetic modifiers (Kapusta & Feschotte, 2014; Necsulea et al, 2014; Hezroni et al, 2015, 2017; Chen et al, 2016; Toiber et al, 2017; Carlevaro-Fita et al, 2019). Significantly, a number of studies have proposed that transposable elements (TEs) that coincide with lncRNAs, referred to as Repeat Insertion Domains of LncRNAs (RIDLs), often constitute the conserved functional domains of these lncRNAs (Kelley & Rinn, 2012; Kapusta et al, 2013; Carlevaro-Fita et al, 2019; Lee et al, 2019; Plaisance et al, 2023).

Additionally, these TEs play a role in determining subcellular localization and can act as transcription regulators by recruiting transcription factors to their loci, giving rise to tissue specificity. However, it’s crucial to note that this correlation doesn’t universally apply, as some lncRNAs deviate from these trends. This variation poses a challenge in comprehensively understanding the characteristics and roles of conserved lncRNAs. Notably, two TE classes, retrotransposons, and simple repeats, have gained attention in deciphering evolutionary transitions from zebrafish to humans (Sotero-Caio et al, 2017; Chang et al, 2022; Wells et al, 2023). These elements undergo diverse modes of evolutionary selection, contributing to mosaic-like genomic patterns. Retrotransposons, especially, have been pivotal in fostering genetic diversity, driving innovation, and enabling adaptation throughout the evolutionary journey(Cordaux & Batzer, 2009). Their dynamic nature influences gene structure and regulation, shaping genetic compositions. Simultaneously, simple repeat elements drive localized evolution, contributing to novel traits and species emergence. Their mutability positions them as essential in understanding species diversification, adaptation, and sequence evolution in syntenic regions over time (Cordaux & Batzer, 2009).

The current study explores the functional signatures inherent in syntenic lncRNAs from humans and zebrafish. Our objective was to unravel the evolutionary dynamics underpinning the divergence of lncRNA sequences while preserving functional significance across species. Leveraging publicly available datasets, we computed the expression profiles and examined the chromatin mark signatures of these syntenic conserved lncRNAs. Furthermore, our investigation extends to elucidating the role of transposable elements (TEs) in molding the sequence variations observed within syntenic conserved lncRNAs across species evolution. Such variations could potentially contribute to genomic innovation and foster the conservation of functional attributes within the locus.

## Results

### Syntenic conserved lncRNA between humans and zebrafish

Syntenic conservation is defined as sets of homologous genes conserved across different species on the same chromosome and in the same order (Fig 1A). Syntenic blocks between humans (hg19) and zebrafish (danRer7) were computed using a web-based application called Synteny Portal at a resolution of 150 kb (Lee et al, 2016). A total of 969 synteny blocks were generated between human and zebrafish genomes. These blocks ranged from 150 kb to 12,700 kb in humans and from 160 kb to 450 kb in zebrafish (Supply Fig 1). Next, we normalized the syntenic block sizes from the genome size of the respected animal as the zebrafish genome is approximately half the size of the human genome. Upon size density distribution analysis of the syntenic blocks, we observed a similar trend suggesting the block size variation could be due to genome expansion in humans (Kolmogorov-Smirnov statistic (D) - 0.1961; p-value = 1.1102e-16). (Fig 1 B) (Lane & Martin, 2010) These synteny blocks contained 2 to 460 protein-coding genes. To identify and examine lncRNA which are syntenically conserved across humans and zebrafish, we considered lncRNA if found in the synteny block and originating from similar genomic loci, irrespective of sequence conservation (Fig 1A). For this purpose, we utilized the ZFLNC database, a comprehensive dataset of annotated lncRNAs in zebrafish. In this database, lncRNAs are considered syntenically conserved if they have more than five anchor points within a 20 k region upstream or downstream of the gene (Hu et al, 2018). Using this approach, we identified 337 genes in zebrafish that produce 526 lncRNAs exhibiting synteny conservation with 457 human lncRNA transcripts. We refer to these as zebrafish: human syntenic lncRNAs (Fig 1C). We additionally applied a filter to remove lncRNAs showing conservation based on Blastn search and also removed lncRNAs which were completely overlapping sense protein coding genes (Supply Fig 2B). These syntenic lncRNAs exhibit comparable patterns in length, exon count, and expression profiles compared to non-syntenic lncRNAs (Supply Fig 2-3).

**Figure 1:**
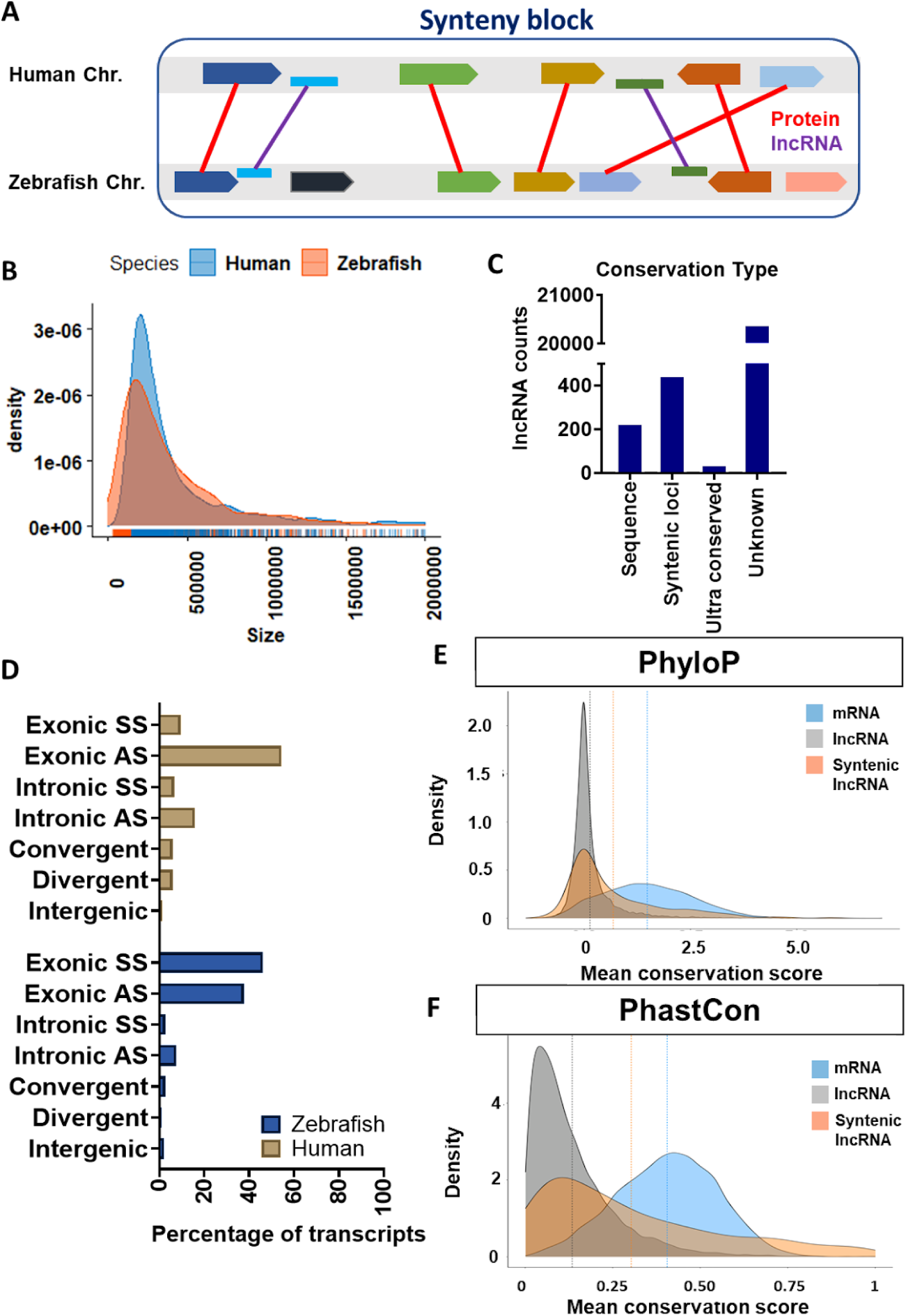
Syntenic lncRNA between zebrafish and humans. [A]Schematic representing the syntenic lncRNAs conservation across human and zebrafish; [B]Density plot of normalized size distribution between conserved syntenic blocks of humans and zebrafish (in bp) (Kolmogorov-Smirnov statistic(D)-0.1961; p-value=1.1102e-16). [C] Distribution of lncRNA based on their mode of conservation in ZFLNC database [D] Genomic position distribution of syntenic conserved lncRNAs in zebrafish and humans. [E,F] PhyloP and Phastcons conservation scores for syntenic lncRNA conserved between human and zebrafish, all lncRNAs and mRNAs from 100 vertebrate different species.

**Figure 2:**
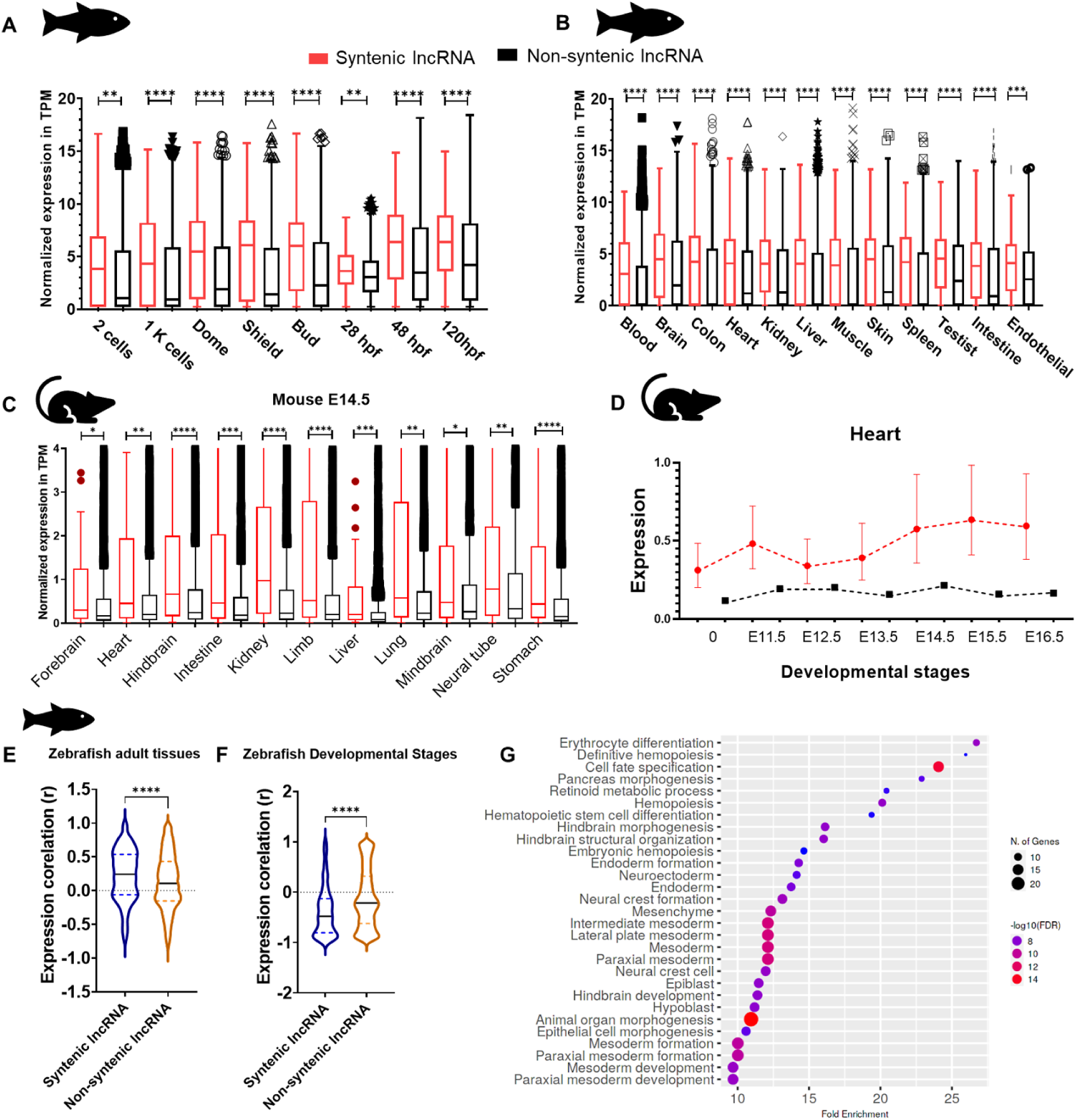
Expression profile of syntenic lncRNAs in zebrafish. [A, B] Expression profile of syntenic and non-syntenic lncRNAs across early development and adult tissues of zebrafish; *** P<0.001,**** P<0.0001 (Mann-Whitney U); [C]Expression profile of syntenic and non-syntenic lncRNAs across early development stage of mouse(E14.5) **** P<0.0001; *** P<0.001; ** P<0.01;* P<0.05 (Mann-Whitney U); [D] Dynamic expression profile of mouse heart tissues represented as geometric mean with 95% CI for syntenic and non-syntenic lncRNAs. [E,F] Expression correlation (r) of syntenic and non-syntenic lncRNAs with its neighboring protein-coding genes during early development and adult tissues of zebrafish; **** P<0.0001 (Mann-Whitney U) [G]The dot plot represents syntenic lncRNA’s neighboring protein-coding gene-associated phenotype ontology. The color represents the -log10 (FDR) value and the circle represents the number of genes associated to the phenotype.

**Figure 3:**
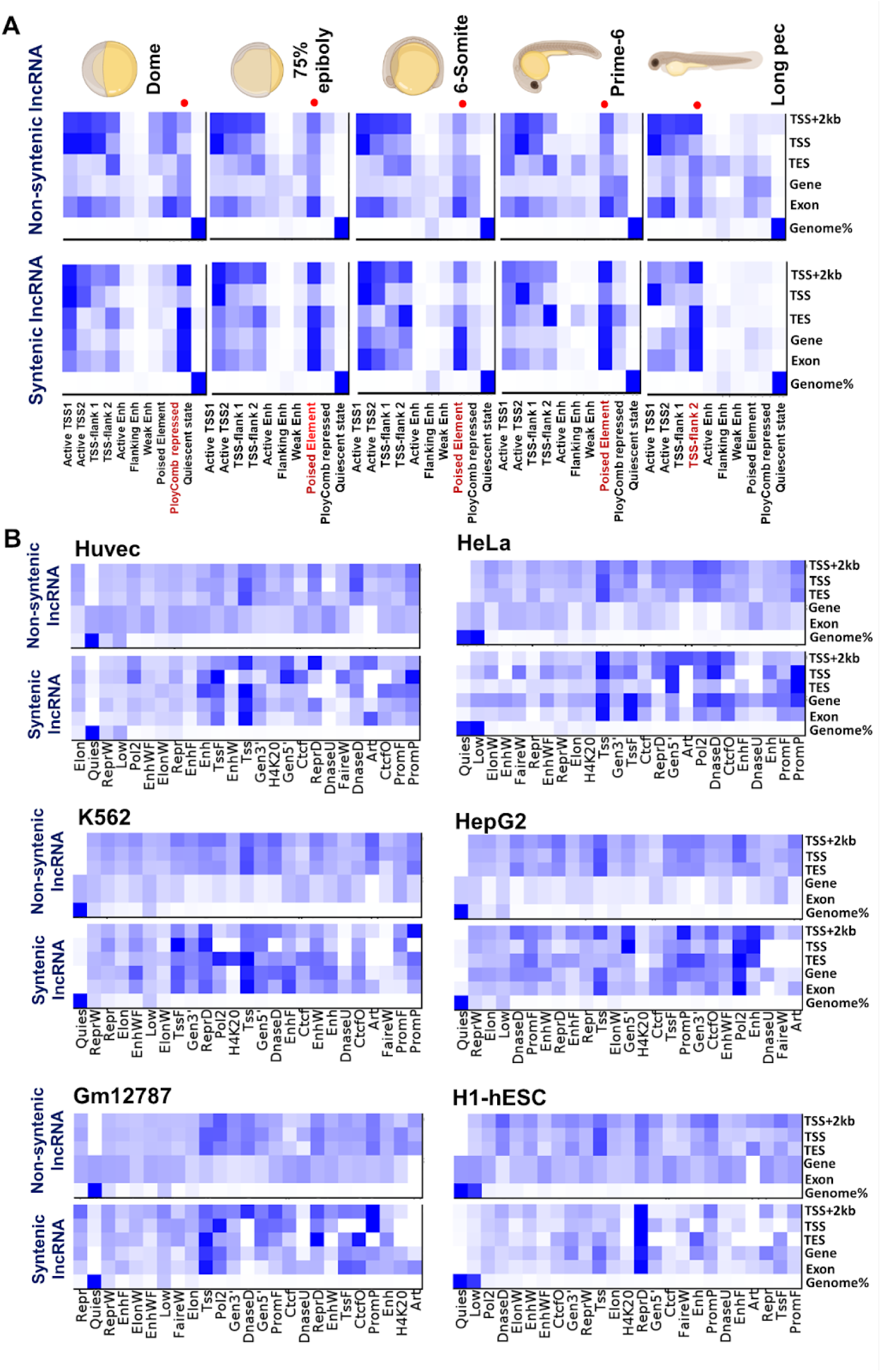
Chromatin state marks overlapping syntenic lncRNAs. [A] Overlap of syntenic and non-syntenic lncRNAs across chromatin states of 5 zebrafish developmental stages. The Chromatin state is represented in the y-axis and the classification of lncRNA is represented in the x-axis. Color represents the enrichment of the chromatin state. Highlighted with red represented chromatin state in focus. The heatmap has column specific color scale wherein white will correspond to 0 and the darkest color will be the maximum value [B] Overlap of syntenic and non-syntenic lncRNAs across chromatin states of 6 different cell lines (HUVEC, HeLa, HepG2, Gm12787, K562, and H1-hESC). The Chromatin state is represented in the y-axis and the classification of lncRNA is represented in the x-axis. Color represents enrichment of the chromatin state;

Next, we categorized the identified zebrafish syntenic lncRNAs according to their genomic positions. We noted that a significant portion of syntenic lncRNAs in zebrafish overlapped with exons, either on the sense strand (162 lncRNA transcripts) or the antisense strand (125 lncRNA transcripts), of nearby protein-coding genes. Similarly, in humans, we observed that a majority of syntenic lncRNAs were found to overlap with protein-coding exons, both in the antisense direction (249 lncRNA transcripts) and the sense direction (46 lncRNA transcripts). Furthermore, a subset of syntenic lncRNAs in humans also exhibited overlap with the antisense strand of intronic regions of protein-coding genes (72 lncRNA transcripts) (Fig 1D). We also looked into other recently published lncRNA data sets of zebrafish and found a majority of lncRNA were overlapping with ZFLNC and a very less considerable number of lncRNAs that were novel but didn’t fall under our criteria of syntenic lncRNA, hence we only used the ZFLNC dataset (Nudelman et al, 2018; Yuan et al, 2019; Banerjee et al, 2021; Sehgal et al, 2021).

As an alternative to ascertain sequence conservation, we calculated average phyloP and phastCons scores for protein coding, all lncRNAs, and syntenic lncRNAs from 100 vertebrates which includes subsets of Primate, Euarchontoglires, Laurasiatheria, Afrotheria, Mammal, Aves, Sarcopterygii and Fish species. The density distribution of the mean conservation scores showed low nucleotide conservation from phyloP and PhastCons score for syntenic lncRNAs (p<0.005; two-tailed Wilcoxon rank sum test) when compared to mRNA and all zebrafish lncRNAs (Siepel et al, 2005; Pollard et al, 2010). (Fig 1E-F).

### Syntenic lncRNAs show an abundance in expression

We performed expression profile analysis of syntenic lncRNAs across different developmental stages (2-4 cells, 1K cells, dome, shield, bud, 28 hours post-fertilization (hpf), 48 hpf, and 5 days post-fertilization (dpf)) and different tissues of adult zebrafish (blood, brain, colon, endothelial, heart, intestine, kidney, liver, muscle, skin, spleen, and testis) by reanalyzing publicly available RNA seq datasets (Pauli et al, 2012; Yang et al, 2020; Sehgal et al, 2021). We observed that syntenic lncRNAs show ∼4-fold increase in mean expression when compared to non-syntenic lncRNAs during early developmental stages from the 2 cell to bud stage. During the late developmental stages, there was a ∼2 fold increase in mean expression when compared to non-syntenic lncRNAs during 28 hpf to 120 hpf (Mann–Whitney test p=<0.0001). In the adult tissues of zebrafish, we see ∼3-fold increase in the mean expression of syntenic lncRNA when compared to non-syntenic lncRNAs in all the tissues analyzed (Fig 2 A-B). We further examined if this pattern is consistent across species by analyzing mouse syntenic lncRNAs. We observed a similar pattern of expression for mouse syntenic conserved lncRNAs in different tissues during early development analyzed using ENCODE data (Supply table 1). We observed a higher expression pattern in syntenic lncRNAs compared to non-syntenic lncRNAs (Fig 2C). Additionally, these syntenic lncRNA exhibit dynamic expression patterns during the development of both mice and zebrafish (Fig 2 D)(Supply Fig 4-5). Increased expression of these syntenic lncRNAs in zebrafish developmental stages and adult tissues suggests they might be abundant during these processes and may be important for proper development and tissue homeostasis in zebrafish.

**Figure 4:**
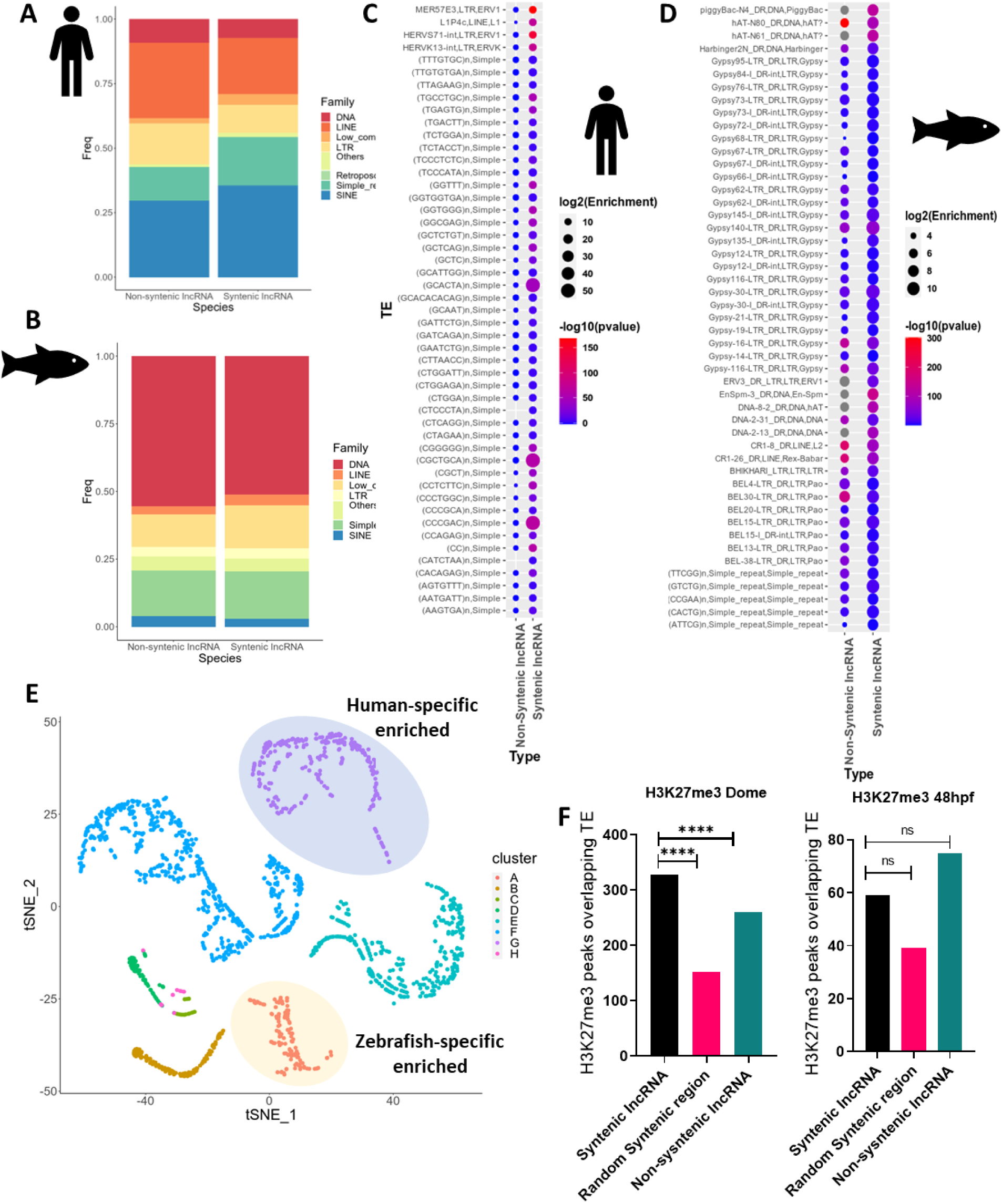
TE contributing to syntenic evolution. [A, B] Stacked bar plot representing percentage distribution of TE family classes in humans and zebrafish overlapping syntenic and non syntenic lncRNAs respectively; [C, D] Dot plot representing enrichment to top selected TEs in humans and zebrafish. The circle represents the log2(enrichment) value and the color represents -log10(p-value). [E] tSNE plot representing 8 clusters of TEs representing conserved enrichment between human and zebrafish. [F] Bar plot representing the overlap between lncRNAs, TEs, and H3K27me3 peaks in syntenic lncRNAs, Random Syntenic regions and non-syntenic lncRNAs in Dome and 48 hpf zebrafish. ns-not significant; **** P<0.0001 (Fisher’s t-Test)

**Figure 5:**
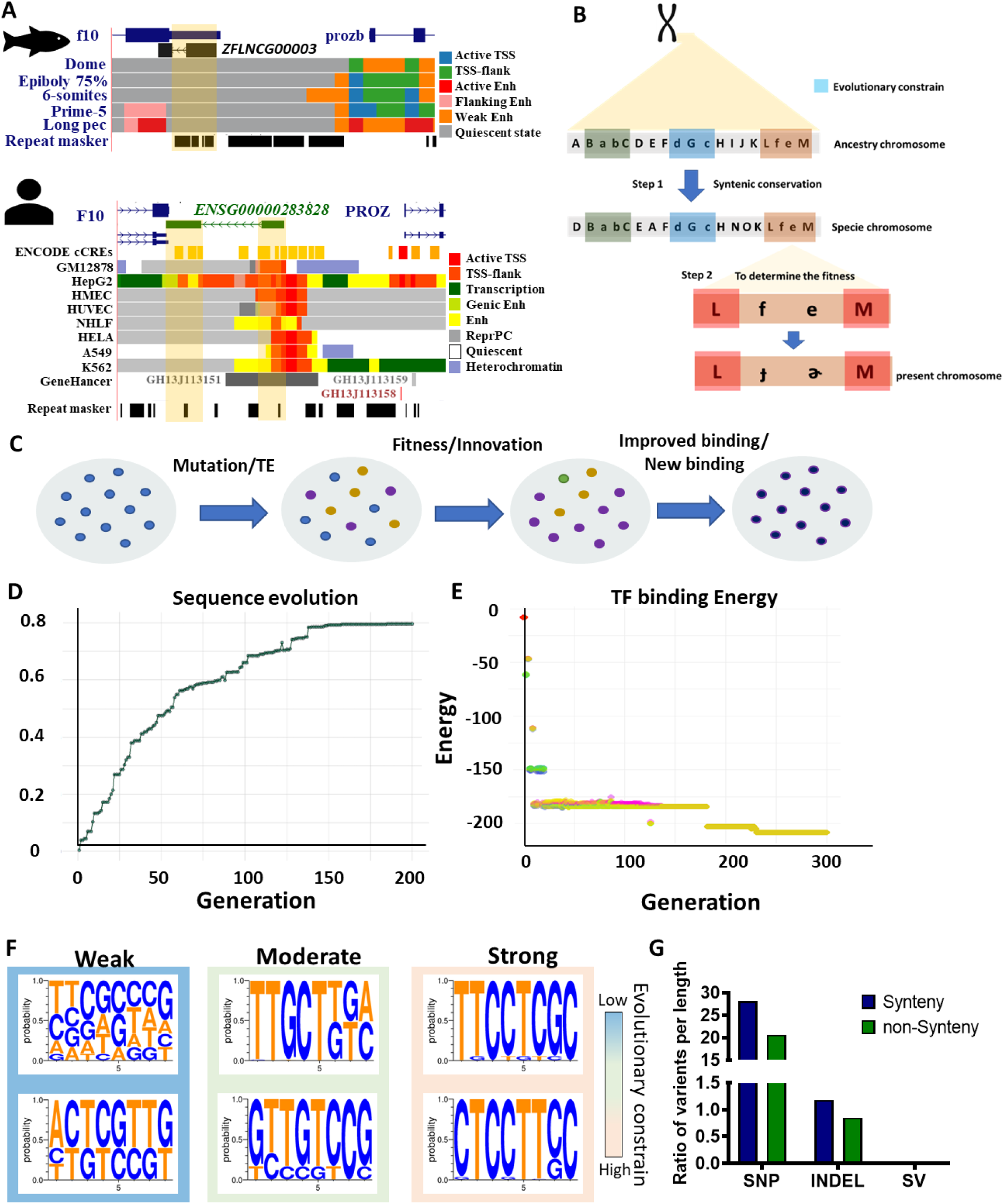
Evolution of the syntenic region is a two-way process. [A] UCSC genome browser screenshot of *ZFLNCG00003* lncRNA locus in zebrafish and *ENSG00000283828* lncRNA locus in human. [B] Illustration of hypothetical model for the two-way evolutionary process of syntenic lncRNA sequence in an organism. The first step is the transfer of syntenic conserved regions from the ancestral genome and the second step is the rapid mutation to attain fitness for survival. [C]Schematic representation of evolution of sequence in an organism mediated by natural selection. [D]Line plot representing genomic simulation of 8kb DNA using genetic algorithm ESTReMo for 1000 generations. The genome got stabilized after 140 generation. The y axis represent the fitness of the organism and the x axis represent the generation [E] Dot plot representing binding energy of TF motif at the y axis across the generation in x axis. [F]Representation of TF binding motif logos across the generation after evolution. [E] Bar plot representing ratio of variant per length in SNP, Indels and SVs from 1000 genome project.

We also analyzed syntenic lncRNA-protein coding gene pairs (530 pairs) and non-syntenic pairs (9776 pairs) in zebrafish developmental stages. Syntenic pairs showed a negative correlation, while non-syntenic pairs did not. In adult zebrafish tissue, a slight positive correlation was observed in both pairs (Mann–Whitney test p=<0.0001) (Fig 2E-F). Gene ontology analysis revealed that protein-coding genes near syntenic lncRNAs are associated with differentiation and morphogenesis (Fig 2G), suggesting a role in early organism development.

### Syntenic lncRNAs are enriched with *cis*-regulatory signatures

Having over ∼ 92% of the zebrafish syntenic lncRNA loci located in proximity to protein-coding genes (Fig 1D), we investigated whether these lncRNA loci contain *cis*-regulatory elements (CREs) or not. We used the *cis*-regulatory element annotation by the Danio-Code consortium which used ChromHMM to annotate the genomic regulatory regions based on 4 major histone signatures (H3K4me3, H3K4me1, H3K27me3, and H3K27ac) (Ernst & Kellis, 2017; Baranasic et al, 2022) and classified them into 10 types (Active TSS 1, Active TSS 2, TSS Flanking region 1, TSS Flanking region 2, Active enhancer 1, Weak enhancers, Primed enhancer, Poised elements, Polycomb repressed regions and Quiescent state) across 5 developmental stages (Dome, 75% epiboly, 5-6 somite, prime5, and long pec stage) (Fig 3A). We performed an analysis to compare the chromatin state annotations overlapping on syntenic lncRNA loci and non-syntenic lncRNA loci across different zebrafish developmental stages. We observed that the syntenic lncRNA loci showed significant enrichment (4.7-fold) of polycomb repression marks during the dome stage. Additionally, we found syntenic lncRNA loci with enrichment (∼5-fold) of poised element marks during the 75% epiboly stage, 5-6 somite stage, and prime5 stage, as well as enrichment (3.4-fold) of TSS Flanking region 2 marks during the long pec stage. These marks were predominantly present on the gene body, exon, transcript end site (TES), and 2kb upstream of the transcription start site (TSS+2kb) of the syntenic lncRNAs in zebrafish. (Fisher T-test, p-value < 0.0001). Next, we examined the enrichment of these chromatin marks on randomly selected, size-matched synthetic regions. It is known that regulatory blocks are abundant within synthetic blocks and could contain regulatory elements like enhancers. Our analysis showed no noticeable enrichment of chromatin signatures in the randomly selected, size-matched syntenic regions (Supply Fig 6). This indicates that the observed enrichment is specific to syntenic lncRNA loci in zebrafish.

Additionally, we delved into chromatin marks in various human cell lines (HUVEC, HeLa, HepG2, Gm12787, K562, and H1-hESC) by utilizing the Epigenome Roadmap Project dataset to compare it to the observed trend found in zebrafish (Fig 3B). Our observations revealed intriguing patterns. In HUVEC, HeLa, K562, and Gm12787, there was notable enrichment of transcription start site signatures within the gene body, exons, and transcript end sites (TES) of syntenic lncRNAs, in contrast to non-syntenic lncRNAs in humans. HepG2 cell lines exhibited significant enrichment of enhancer and Pol2 signatures in and around the syntenic lncRNA locus compared to non-syntenic lncRNAs. Particularly interesting was the presence of repressor marks in H1-hESC, predominantly on the gene body, exons, TES, and 2kb upstream of the transcription start site (TSS+2kb) of syntenic lncRNAs, as opposed to non-syntenic counterparts. Notably, these findings correlated with zebrafish’s early developmental stages, suggesting that a substantial subset of these syntenic lncRNAs plays a crucial role in modulating target gene expression during embryonic development.

Next, we scanned the syntenic lncRNA loci for transcription factor (TF) binding, by overlapping the syntenic lncRNA loci with the JASPER 2022 TF database for zebrafish (Castro-Mondragon et al, 2022). Upon analyzing the gene ontology for molecular and biological processes for TF which overlapped the syntenic lncRNAs, we observed that these TF were *cis*-regulating and mainly involved in cell fate specification and embryonic organ development and morphogenesis (Supply Fig 7). This enrichment of chromatin marks (polycomb repression, poised element, and TSS Flanking region 2) across developmental stages, and correlated expression with their neighboring protein-coding genes during development and in adult tissues respectively suggests that syntenic lncRNA loci could plausibly act as *cis*-regulatory elements controlling the neighboring genes expressions during cell fate specification and embryonic organ development and morphogenesis in zebrafish(Supply Fig 8).

### Transposable elements mediated evolution of conserved syntenic lncRNAs

We investigated factors influencing sequence variations in lncRNA loci across species. Rapid evolution in these regions, driven by low evolutionary constraints, is a key factor. Transposable elements (TEs) also play a significant role in sequence expansion and evolution. Analyzing TE overlaps with syntenic lncRNAs, we found differences in TE family contributions between humans and zebrafish. In humans, SINE and simple repeat elements are elevated in syntenic lncRNAs, while in zebrafish, LTR family elements show an increase. Notably, zebrafish exhibit a substantial increase in DNA elements and a decrease in SINE elements compared to humans (Fig 4A-B). Enrichment analysis shows unique TE classes in human and zebrafish syntenic lncRNAs. In humans, there’s an enrichment of simple repeat sequences, while in zebrafish, Gypsy, BEL/Pao, and hAT classes are enriched (Fig 4 C-D). Interestingly, the BEL/Pao class is absent in mammals but present in metazoans, influencing sequence evolution (de la Chaux & Wagner, 2011). We generated a t-SNE plot of TE enrichment data from humans and zebrafish, revealing eight clusters (Fig 4E). Cluster G shows TEs enriched in syntenic lncRNAs in humans, with some in zebrafish, mainly simple repeats (36.5%) and LTR repeats (21.9%). Cluster A is enriched in zebrafish, mostly with direct repeat elements (78.1%), with a minor presence in some human TEs. Other clusters display minimal enrichments specific to humans or zebrafish (Fig 4E). Overall, species-specific TEs contribute to sequence variation in noncoding regions.

TEs are known for their role in regulating the transcription of nearby protein-coding genes. To confirm our observations during zebrafish developmental stages and investigate the involvement of TEs, we conducted an analysis to examine the overlap of H3K27me3 peaks (de la Calle Mustienes et al, 2015) on TEs that coincide with syntenic lncRNAs (531 lncRNA transcripts), size-matched random syntenic regions (531 regions), and randomly selected non-syntenic lncRNAs (531 lncRNA transcripts). In the dome developmental stage of zebrafish, our findings indicate a significant enrichment of H3K27me3, a repressor mark, within syntenic lncRNAs overlapping TEs compared to size-matched random syntenic regions (Fisher’s exact test; p-value = 7.013e-16; 95% confidence interval [CI]: 1.778080 to 2.641569; Odds Ratio [OR] - 2.16317). Similarly, significant enrichment was observed when comparing syntenic lncRNAs overlapping TEs to non-syntenic lncRNAs overlapping TEs (Fisher’s exact test; p-value = 0.006159; 95% CI: 1.066794 to 1.490184; OR - 1.260346). Conversely, during the 48 hpf developmental stage of zebrafish, no significant H3K27me3 enrichment was observed in syntenic lncRNAs overlapping TEs compared to size-matched random syntenic regions overlapping TEs (p-value = 0.05348; 95% CI: 0.9941693 to 2.3395438; OR - 1.517508), nor when comparing syntenic lncRNAs overlapping TEs to non-syntenic lncRNAs overlapping TEs (Fisher’s exact test; p-value = 0.1925; 95% CI: 0.5472433 to 1.1206871; OR - 0.7847096). These results are consistent with the observed chromatin state overlap enrichment during the dome and long pec stages of zebrafish development (Fig 4F)(Supply Fig 9). Specifically, in the early developmental stage (dome), there was an enrichment of repressive marks on syntenic lncRNAs, which transitioned to enhancer or TSS marks later in the developmental stage. These insights underscore the dynamic regulatory role of syntenic lncRNAs, TEs, and their impact on chromatin states, contributing to the intricate regulatory landscape during zebrafish development.

### Conserved *cis*-regulation with sequence robustness and innovation at syntenic lncRNA loci

Using an illustrative case of a syntenic conserved lncRNA *ENSG00000283828* in human and *ZFLNCG00003* lncRNA in zebrafish, we next aimed to discern the underlying factors contributing to the conservation of syntenic regions regardless of sequence (Fig 5A). Notably, our analysis revealed distinct features in exon 1 of both *ZFLNCG00003* lncRNA in zebrafish and *ENSG00000283828* lncRNA in humans. In zebrafish, exon 1 exhibited the presence of TEs AGEL from the hAT family, U1AR_DR from the dada family and (TATTAA)n from the simple repeat family. In contrast, the equivalent region in humans harbored the L1NB4 element of the L1 family, while exon 2 contained MER5B from the hAT family. Additionally, we noted the occurrence of TEs in the intronic and promoter regions of these lncRNAs, indicating the potential involvement of TE-mediated mechanisms in driving sequence evolution. Based on earlier findings indicating the functional relevance of DNA elements within the lncRNA locus, our investigation focused on the transcription factor (TF) binding to this region in both humans and zebrafish. Intriguingly, our analysis unveiled a notable disparity in TF binding motifs between the human and zebrafish lncRNA loci. Specifically, the human *ENSG00000283828* lncRNA locus exhibited a considerably greater number of significant TF binding motifs compared to its zebrafish counterpart *ZFLNCG00003* lncRNA. Moreover, the overlap or conservation of TF binding motifs between the two species was minimal, implying a potential emergence of this locus as a vertebrate innovation due to sequence evolution (Supply Fig 10).

We hypothesize that syntenic region evolution is a two-step process as depicted in Fig 5B. Initially, the syntenic regions or blocks in the genomes are transferred to the next species in the same order and on the same chromosome. This implies the first step in evolution. After that, the species undergoes mutational selection to increase its fitness for survival, resulting in a change in sequences across the genome. These changes are major in the non-coding regions of the genome because the evolutionary constraints on the protein-coding genes are high compared to non-coding genes (Fig 5C). Next, we used an evolutionary simulator of transcription regulatory networks (ESTReMo), a genetic machine learning algorithm to understand the transcription factor binding motif evolution. We observed that in our candidate, there was no sequence conservation, yet 451 TF-binding proteins were found to be binding to the loci in humans and zebrafish. This is due to the sequence robustness of the TF binding protein or the addition of new TF binding motifs in the regions that have been previously known. To understand this phenomenon we simulated an 8 kb DNA region from the zebrafish genome considering it as an organism with parameters indicating the region to harbor 8bp as TF motif regions and simulation (evolving) till 1000 generation. It was observed that upon simulation the region incorporated mutation in the zebrafish lncRNA locus which led to an increase in the fitness of the system and after 140 generations the saturation was achieved for the system with maximum fitness (Fig 5D).

We next observed the binding efficiency of the TF binding motifs. We found that with the increasing number of generations mutations caused in the TF binding motifs led to the binding energy of the motifs being increased subsequently with each passing generation (Fig 5E). When we looked into the sequences of the motifs we found that 3 types of motif evolution had occurred in this locus. The first type of motif evolution indicated that there was a drastic change in the sequence of the motif to attain higher binding energy which indicates that these types of motifs exhibit low evolutionary constraints. The second type of motif evolution indicated moderate changes in the sequence of the motif to exhibit higher binding energy indicating moderate evolutionary constraints on these motifs. The last type of motif evolution implies that there are motif regions with high evolutionary constraints that exhibit no or very low sequence chances to attain higher binding efficiency of the TF (Fig 5F).

To further check the hypothesis we looked into the mutational landscape across the human genome using the 1000 genome project dataset and computed the frequency of mutations in the syntenic and non-syntenic regions. We observed that the ratio of the number of variants per length in both the syntenic and non-syntenic regions led to the conclusion that there is an increase in the variation in the syntenic region. This data suggests that the mutations accusing the organism are higher in syntenic regions compared to the non-syntenic regions (Fig 5G). Overall this study indicates that syntenic conserved lncRNA across species are enriched with *cis*-regulatory regions and the sequence divergence observed between human and zebrafish is due to the rapid evolution of sequence at their TF binding regions which are robust to binding of these TF.

## Discussion

Long noncoding RNA is known to be poorly sequence conserved, which elicits the need to investigate other alternative modes of evolutionary conservation. In this study, we probed into the syntenic mode of evolutionary conservation of lncRNA in humans and zebrafish. It was observed that a large proposition of syntenic lncRNA was found to be sense or antisense overlapping to protein-coding exonic regions and in very close proximity to their neighboring protein-coding genes. This observation might be influenced by bias, as syntenic block annotations primarily rely on protein-coding genes as anchors or reference points, potentially leading to a higher likelihood of protein-coding genes overlapping with sense or antisense lncRNAs. Previous studies have shown that multiple lncRNA which were in proximity to neighboring protein-coding genes were known to be *cis-*regulating their expression (Gil & Ulitsky, 2020). This cis-regulation was either due to the escorting of TFs to the regulatory regions by the RNA molecules or the presence of DNA regulatory elements on the lncRNA loci (Yap et al, 2010; Schertzer et al, 2019; Chouvardas et al, 2022; Winkler et al, 2022). This compelled us to also look into whether the syntenic lncRNA exhibits a cis-regulatory signature that governs the expression of the neighboring genes. We observed that syntenic lncRNA was enriched with regulatory marks on gene body, exons, TES, and promoter regions of lncRNA when compared to non-syntenic lncRNA using chromHMM chromatin state annotation of zebrafish. These regulatory regions were observed to be in the suppressed or poised state during early development (dome, 75% epiboly, and prime 5) and later activated during the long pec stage. A similar pattern emerged in human cell culture datasets, where H1-hESC cells demonstrated trends akin to the early developmental stage of zebrafish. Conversely, other cell types exhibited context-specific functionalities, aligning with the understanding that lncRNAs often exhibit tissue or cell type-specific roles (Ulitsky et al, 2011; Derrien et al, 2012; Pauli et al, 2012; Kaushik et al, 2013; Haque et al, 2014; Akay et al, 2019; Mattioli et al, 2019). These findings are consistent with previously analyzed data on antisense lncRNAs, which exhibited comparable correlation trends in zebrafish (Pillay et al, 2021). It is intriguing to consider conducting functional studies to gain insights into their molecular roles. Notably, past knockout studies of lncRNAs selected for functional role in zebrafish have underscored that these molecules might not overtly impact embryogenesis, viability, and fertility but could have roles that may be redundant, subtle, or context-dependent.(Goudarzi et al, 2019). Altogether the comprehensive analysis of expression, correlation, and chromatin enrichment data collectively suggests that syntenic lncRNAs confer DNA regulatory functions as one of their functional modalities. Through this mechanism, they fine-tune cis-regulatory mechanisms crucial for organismal development. The role of other functional modalities cannot be ruled out, as multiple studies have suggested that a lncRNA could possess more than one functional module. lncRNAs

The evolutionary trajectory of syntenic lncRNA sequences has emerged as a captivating area of interest. Multiple mechanisms underpin the sequence alterations in noncoding regions, driven by relaxed evolutionary constraints (Derrien et al, 2012; Kapusta & Feschotte, 2014; Hezroni et al, 2015). Transposable elements (TEs) have played a significant role in the expansion of noncoding genomic regions, impacting transcription regulation and subcellular localization, with some TEs notably implicated in modulating lncRNAs(Kelley & Rinn, 2012; Kapusta et al, 2013; Johnson & Guigó, 2014; Carlevaro-Fita et al, 2019; Plaisance et al, 2023). Interestingly, a distinct subset of TE families exhibited enrichment in both humans and zebrafish. Particularly in zebrafish, the Gypsy class and BEL/Pao elements from the LTR family, which are significant retrotransposons associated with genome evolution and rearrangement, exhibited notable enrichment. Notably, the absence of BEL/Pao elements in humans highlights an intriguing observation and potential function of these TEs in zebrafish (de la Chaux & Wagner, 2011; Bai et al, 2021; Chang et al, 2022). These LTRs have also been linked to or implicated in zygotic genome activation in zebrafish (Chang et al, 2022). In contrast, human syntenic lncRNAs exhibited substantial enrichment of simple repeat TEs. These simple repeat elements are recognized contributors to genome expansion and have demonstrated roles in local or adaptive evolution (Kashi & King, 2006; Yuan et al, 2021). Their expansion is known to introduce new transcription factor binding motifs or regions conducive to epigenetic regulation and genomic imprinting, potentially leading to vertebrate innovation of various phenotypes and functions(Sutherland & Richards, 1995; Katti et al, 2001; Kashi & King, 2006; Yuan et al, 2021). While the conservation of cis-regulatory functions and functional modalities, such as DNA elements within the loci of these syntenic lncRNAs, are shared mechanisms in both humans and zebrafish, the rapid evolution of their sequences continues to spark interest. It’s plausible that the expansion and evolution facilitated by transposable elements (TEs) could contribute to vertebrate innovation(Etchegaray et al, 2021; Almeida et al, 2022; Modzelewski et al, 2022). This might offer an explanation for the observed sequence variations among these lncRNAs despite their conservation based on syntenic positioning across species.

In this context, we hypothesize a two-step evolutionary process. Initially, syntenic regions are transferred to the next species while retaining their order in chromosomes. Subsequently, mutations occur to enhance the fitness of these syntenic lncRNAs, which function via the DNA elements on their locus, driving selective sequence changes and increasing the binding fitness of TFs across these regions. The binding motifs of transcription factors at enhancers and promoters exhibit sequence robustness and have evolved across the species while retaining the functionality (Payne & Wagner, 2014; Pérez-Rico et al, 2020). Regulatory regions such as enhancers don’t strictly demand the exact sequence conservation seen in evolution, indicating that their sequence constraint might stem from additional regulatory or functional roles (Snetkova et al, 2021). These evolutionary processes may also extend to other regulatory regions within the syntenic block, although additional detailed analyses are necessary to elucidate this possibility fully. Furthermore, it’s important to acknowledge that other evolutionary mechanisms may also contribute to the sequence divergence of syntenic lncRNA loci.

These findings collectively underscore the significance of syntenic conserved lncRNAs shared between humans and zebrafish. These lncRNAs exhibit enrichment of cis-regulatory functions and are predominantly located in proximity to protein-coding genes, either in sense or antisense orientation. The analysis of expression and correlations, coupled with the enrichment of cis-regulatory signatures, implies that these syntenic lncRNAs are primarily involved in repressive roles during the early developmental stages of organisms, thereby influencing the regulation of their cis-associated protein-coding genes involved in development. In light of our findings, our two-way evolution hypothesis attempts to elucidate one of the underlying reasons for the observed sequence variations within these syntenic conserved lncRNAs across species.

## Methods

### Syntenic Blocks between humans and zebrafish

We used an online tool called synteny portal (http://bioinfo.konkuk.ac.kr/synteny_portal/htdocs/synteny_builder.php) to generate synteny conserved blocks between human and zebrafish at a resolution of 150kb. The conserved syntenic block and their circos plot were downloaded from the portal. The density map of the length of the syntenic blocks was plotted after normalizing it with the size of the individual genome using R ggplot function.

### Genomic position annotation of syntenic lncRNA

We used a bespoke script with sight modification from github (https://github.com/gold-lab/shared_scripts/tree/master/lncRNA.annotator) for lncRNA positional annotation. The annotation classifies the lncRNAs into unstranded, intergenic; samestrand (SS), lincRNA upstream; divergent; samestrand, protcod upstream; convergent; intronic_Antisense (AS); intronic_SS; exonic_AS; exonic_SS; and unstranded, genic based on their genomic position.

### Syntenic lncRNA conservation based on PhyloP and Phastcon score

The PhyloP and Phastcon score (from 100 vertebrate species) for the whole human genome was downloaded from USCS table browser (Siepel et al, 2005; Pollard et al, 2010). The score overlapped with the syntenic lncRNA, all lncRNA and the mRNA position of conserved genes using bedtools and the density plot was plotted using R(Quinlan & Hall, 2010).

### Expression analysis of different developmental stages and adult tissues of zebrafish

We performed RNA seq analysis of the previously published data sets (Pauli et al, 2012; Yang et al, 2020; Sehgal et al, 2021). We generated a custom genome annotation file (GTF file) by merging the GTF annotation of lncRNA from the ZFLNC database to the reference ensemble annotation GTF (Hu et al, 2018). Quality check was performed using FastQC (https://www.bioinformatics.babraham.ac.uk/projects/fastqc/) and reads were trimmed using trimmomattic (Bolger et al, 2014). The genomic alignment was performed using STAR v2.7 and raw counts were obtained using the htseq-count function with the intersection-strict option (Dobin et al, 2013; Anders et al, 2015). The data were normalized (TPM) using the DESeq2 normalization function in R studio and plotted using GraphPad or R (Love et al, 2014). All the lncRNA without any expression in any of the tissue or developmental stages were removed from the analysis.

### Expression correlation between lncRNA-neighboring protein-coding genes in different developmental stages and adult tissues of zebrafish

We identified the neighboring protein coding genes of lncRNA using bedtools closest function and made pairs for syntenic and non-syntenic lncRNAs with their neighboring protein-coding genes. We took the expression values of the lncRNA and their neighboring protein-coding gene from above analyzed data. Using R we computed the expression Pearson correlation between the lncRNA and their neighboring protein-coding genes and plotted the correlation value using Graphpad v9.

### Bioinformatic analysis of cis-regulatory elements on lncRNAs during zebrafish developmental stages

To identify cis-regulatory elements overlap with syntenic lncRNAs we used previously annotated 10 major chromatin marks for 5 different developmental stages of zebrafish from the Dani Code consortium (Tan et al, 2016; Baranasic et al, 2022) and 6 human cell line data was downloaded from the Epigenome roadmap project (Roadmap Epigenomics Consortium et al, 2015). We used ChromHMM overlap enrichment function with default parameter to compute the overlap of syntenic lncRNAs and non syntenic lncRNA with chromatin annotation of developmental stages (Ernst & Kellis, 2012). Fisher’s t-test was used to compute statistical significance between syntenic lncRNA enrichment and non-syntenic lncRNA.

### TE enrichment of syntenic lncRNA

The repeat masker data for humans (hg38) and zebrafish (zv9) were downloaded from UCSC (Smit, 1996, 1999) and overlapped with syntenic lncRNAs of humans and zebrafish respectively. The enrichment was calculated as previously described (Karakülah & Suner, 2017) and Fisher’s test was computed for the p-value and odds ratio using R.

### Simulation of genome

Simulation of the DNA was performed as described previously using an evolutionary simulator of transcription regulatory networks (ESTReMo) algorithm(O’Neill et al, 2014). Random 8 kb DNA region from the zebrafish genome was taken as an input fasta for simulation with a population size of 500 and generation count of 10000. The promoter length was defined as 128bp based on the average length of the promoter in zebrafish and 8 bp bin size for the TF binding motif. The mutation rate for the promoter was defined as 0.001 and for a motif, it was defined as 0.005. The simulation was performed on the Linux Ubuntu system.

## Data Availability

All the data has been provided in a supplementary file.

## Funding

No funding to disclose

## Conflict of Interest Disclosure

The author declares no conflict of interest

## Acknowledgments

The authors would like to acknowledge Mercy Rophina and Arvind Kumar for their input and suggestions in the analysis.

## Author’s contribution

GR, VS, and SS conceived and designed the experiment, GR performed the analysis. GR,VS, and SS analyzed the results. GR wrote the manuscript and GR, VS, and SS edited the manuscript.

## Notes

### Competing Interest Statement

The authors have declared no competing interest.

## Reference

Akay A, Jordan D, Navarro IC, Wrzesinski T, Ponting CP, Miska EA, Haerty W (2019) Identification of functional long non-coding RNAs in C. elegans. BMC Biol 17: 14.

Alexanian M, Maric D, Jenkinson SP, Mina M, Friedman CE, Ting C-C, Micheletti R, Plaisance I, Nemir M, Maison D et al (2017) A transcribed enhancer dictates mesendoderm specification in pluripotency. Nat Commun 8: 1806.

Ali T, Grote P (2020) Beyond the RNA-dependent function of LncRNA genes. Elife 9. doi:10.7554/eLife.60583.

Almeida MV, Vernaz G, Putman ALK, Miska EA (2022) Taming transposable elements in vertebrates: from epigenetic silencing to domestication. Trends Genet 38: 529–553.

Andergassen D, Muckenhuber M, Bammer PC, Kulinski TM, Theussl H-C, Shimizu T, Penninger JM, Pauler FM, Hudson QJ (2019) The Airn lncRNA does not require any DNA elements within its locus to silence distant imprinted genes. PLoS Genet 15: e1008268.

Anders S, Pyl PT, Huber W (2015) HTSeq--a Python framework to work with high-throughput sequencing data. Bioinformatics 31: 166–169.

Bai J, Yang Z-Z, Li H, Hong Y, Fan D-D, Lin A-F, Xiang L-X, Shao J-Z (2021) Genome-Wide Characterization of Zebrafish Endogenous Retroviruses Reveals Unexpected Diversity in Genetic Organizations and Functional Potentials. Microbiol Spectr 9: e0225421.

Banerjee B, Koner D, Karasik D, Saha N (2021) Genome-wide identification of novel long non-coding RNAs and their possible roles in hypoxic zebrafish brain. Genomics 113: 29–43.

Baranasic D, Hörtenhuber M, Balwierz PJ, Zehnder T, Mukarram AK, Nepal C, Várnai C, Hadzhiev Y, Jimenez-Gonzalez A, Li N et al (2022) Multiomic atlas with functional stratification and developmental dynamics of zebrafish cis-regulatory elements. Nat Genet 54: 1037–1050.

Bolger AM, Lohse M, Usadel B (2014) Trimmomatic: a flexible trimmer for Illumina sequence data. Bioinformatics 30: 2114–2120.

de la Calle Mustienes E, Gómez-Skarmeta JL, Bogdanović O (2015) Genome-wide epigenetic cross-talk between DNA methylation and H3K27me3 in zebrafish embryos. Genom Data 6: 7–9.

Carlevaro-Fita J, Polidori T, Das M, Navarro C, Zoller TI, Johnson R (2019) Ancient exapted transposable elements promote nuclear enrichment of human long noncoding RNAs. Genome Res 29: 208–222.

Castro-Mondragon JA, Riudavets-Puig R, Rauluseviciute I, Lemma RB, Turchi L, Blanc-Mathieu R, Lucas J, Boddie P, Khan A, Manosalva Pérez N et al (2022) JASPAR 2022: the 9th release of the open-access database of transcription factor binding profiles. Nucleic Acids Res 50: D165–D173.

Chang N-C, Rovira Q, Wells J, Feschotte C, Vaquerizas JM (2022) Zebrafish transposable elements show extensive diversification in age, genomic distribution, and developmental expression. Genome Res 32: 1408–1423.

de la Chaux N, Wagner A (2011) BEL/Pao retrotransposons in metazoan genomes. BMC Evol Biol 11: 154.

Chen J, Shishkin AA, Zhu X, Kadri S, Maza I, Guttman M, Hanna JH, Regev A, Garber M (2016) Evolutionary analysis across mammals reveals distinct classes of long non-coding RNAs. Genome Biol 17: 19.

Chen W, Zhang X, Li J, Huang S, Xiang S, Hu X, Liu C (2018) Comprehensive analysis of coding-lncRNA gene co-expression network uncovers conserved functional lncRNAs in zebrafish. BMC Genomics 19: 112.

Cho SW, Xu J, Sun R, Mumbach MR, Carter AC, Chen YG, Yost KE, Kim J, He J, Nevins SA et al (2018) Promoter of lncRNA Gene PVT1 Is a Tumor-Suppressor DNA Boundary Element. Cell 173: 1398–1412.e22.

Chouvardas P, Dhaka B, Zimmerli M, Hanhart D, Moser M, Guillen-Ramirez H, Mishra S, Esposito R, Polidori T, Widmer M et al (2022) Functional identification of cis-regulatory long noncoding RNAs at controlled false-discovery rates.

Cordaux R, Batzer MA (2009) The impact of retrotransposons on human genome evolution. Nat Rev Genet 10: 691–703.

Derrien T, Johnson R, Bussotti G, Tanzer A, Djebali S, Tilgner H, Guernec G, Martin D, Merkel A, Knowles DG et al (2012) The GENCODE v7 catalog of human long noncoding RNAs: analysis of their gene structure, evolution, and expression. Genome Res 22: 1775–1789.

Diederichs S (2014) The four dimensions of noncoding RNA conservation. Trends Genet 30: 121–123.

Dobin A, Davis CA, Schlesinger F, Drenkow J, Zaleski C, Jha S, Batut P, Chaisson M, Gingeras TR (2013) STAR: ultrafast universal RNA-seq aligner. Bioinformatics 29: 15–21.

Ernst J, Kellis M (2012) ChromHMM: automating chromatin-state discovery and characterization. Nat Methods 9: 215–216.

Ernst J, Kellis M (2017) Chromatin-state discovery and genome annotation with ChromHMM. Nat Protoc 12: 2478–2492.

Etchegaray E, Naville M, Volff J-N, Haftek-Terreau Z (2021) Transposable element-derived sequences in vertebrate development. Mob DNA 12: 1.

Furlan G, Gutierrez Hernandez N, Huret C, Galupa R, van Bemmel JG, Romito A, Heard E, Morey C, Rougeulle C (2018) The Ftx Noncoding Locus Controls X Chromosome Inactivation Independently of Its RNA Products. Mol Cell 70: 462–472.e8.

Gil N, Perry RB-T, Mukamel Z, Tuck A, Bühler M, Ulitsky I (2023) Complex regulation of Eomes levels mediated through distinct functional features of the Meteor long non-coding RNA locus. Cell Rep 42: 112569.

Gil N, Ulitsky I (2020) Regulation of gene expression by cis-acting long non-coding RNAs. Nat Rev Genet 21: 102–117.

Goudarzi M, Berg K, Pieper LM, Schier AF (2019) Individual long non-coding RNAs have no overt functions in zebrafish embryogenesis, viability and fertility. Elife 8. doi:10.7554/eLife.40815.

Haque S, Kaushik K, Leonard VE, Kapoor S, Sivadas A, Joshi A, Scaria V, Sivasubbu S (2014) Short stories on zebrafish long noncoding RNAs. Zebrafish 11: 499–508.

Hezroni H, Ben-Tov Perry R, Meir Z, Housman G, Lubelsky Y, Ulitsky I (2017) A subset of conserved mammalian long non-coding RNAs are fossils of ancestral protein-coding genes. Genome Biol 18: 162.

Hezroni H, Koppstein D, Schwartz MG, Avrutin A, Bartel DP, Ulitsky I (2015) Principles of long noncoding RNA evolution derived from direct comparison of transcriptomes in 17 species. Cell Rep 11: 1110–1122.

Howe K, Clark MD, Torroja CF, Torrance J, Berthelot C, Muffato M, Collins JE, Humphray S, McLaren K, Matthews L et al (2013) The zebrafish reference genome sequence and its relationship to the human genome. Nature 496: 498–503.

Hu X, Chen W, Li J, Huang S, Xu X, Zhang X, Xiang S, Liu C (2018) ZFLNC: a comprehensive and well-annotated database for zebrafish lncRNA. Database 2018. doi:10.1093/database/bay114.

Johnson R, Guigó R (2014) The RIDL hypothesis: transposable elements as functional domains of long noncoding RNAs. RNA 20: 959–976.

Kapusta A, Feschotte C (2014) Volatile evolution of long noncoding RNA repertoires: mechanisms and biological implications. Trends Genet 30: 439–452.

Kapusta A, Kronenberg Z, Lynch VJ, Zhuo X, Ramsay L, Bourque G, Yandell M, Feschotte C (2013) Transposable elements are major contributors to the origin, diversification, and regulation of vertebrate long noncoding RNAs. PLoS Genet 9: e1003470.

Karakülah G, Suner A (2017) PlanTEnrichment: A tool for enrichment analysis of transposable elements in plants. Genomics 109: 336–340.

Kashi Y, King DG (2006) Simple sequence repeats as advantageous mutators in evolution. Trends Genet 22: 253–259.

Katti MV, Ranjekar PK, Gupta VS (2001) Differential distribution of simple sequence repeats in eukaryotic genome sequences. Mol Biol Evol 18: 1161–1167.

Kaushik K, Leonard VE, Kv S, Lalwani MK, Jalali S, Patowary A, Joshi A, Scaria V, Sivasubbu S (2013) Dynamic expression of long non-coding RNAs (lncRNAs) in adult zebrafish. PLoS One 8: e83616.

Kelley D, Rinn J (2012) Transposable elements reveal a stem cell-specific class of long noncoding RNAs. Genome Biol 13: R107.

Kopp F, Mendell JT (2018) Functional Classification and Experimental Dissection of Long Noncoding RNAs. Cell 172: 393–407.

Kurian L, Aguirre A, Sancho-Martinez I, Benner C, Hishida T, Nguyen TB, Reddy P, Nivet E, Krause MN, Nelles DA et al (2015) Identification of novel long noncoding RNAs underlying vertebrate cardiovascular development. Circulation 131: 1278–1290.

Lane N, Martin W (2010) The energetics of genome complexity. Nature 467: 929–934.

Latos PA, Pauler FM, Koerner MV, Şenergin HB, Hudson QJ, Stocsits RR, Allhoff W, Stricker SH, Klement RM, Warczok KE et al (2012) Airn transcriptional overlap, but not its lncRNA products, induces imprinted Igf2r silencing. Science 338: 1469–1472.

Lee H, Zhang Z, Krause HM (2019) Long Noncoding RNAs and Repetitive Elements: Junk or Intimate Evolutionary Partners? Trends Genet 35: 892–902.

Lee J, Hong W-Y, Cho M, Sim M, Lee D, Ko Y, Kim J (2016) Synteny Portal: a web-based application portal for synteny block analysis. Nucleic Acids Res 44: W35–W40.

Lewandowski JP, Dumbović G, Watson AR, Hwang T, Jacobs-Palmer E, Chang N, Much C, Turner KM, Kirby C, Rubinstein ND et al (2020) The Tug1 lncRNA locus is essential for male fertility. Genome Biol 21: 237.

Li K, Blum Y, Verma A, Liu Z, Pramanik K, Leigh NR, Chun CZ, Samant GV, Zhao B, Garnaas MK et al (2010) A noncoding antisense RNA in tie-1 locus regulates tie-1 function in vivo. Blood 115: 133–139.

Liu D, Hunt M, Tsai IJ (2018) Inferring synteny between genome assemblies: a systematic evaluation. BMC Bioinformatics 19: 26.

Love MI, Huber W, Anders S (2014) Moderated estimation of fold change and dispersion for RNA-seq data with DESeq2. Genome Biol 15: 550.

Mattick JS, Amaral PP, Carninci P, Carpenter S, Chang HY, Chen L-L, Chen R, Dean C, Dinger ME, Fitzgerald KA et al (2023) Long non-coding RNAs: definitions, functions, challenges and recommendations. Nat Rev Mol Cell Biol 24: 430–447.

Mattioli K, Volders P-J, Gerhardinger C, Lee JC, Maass PG, Melé M, Rinn JL (2019) High-throughput functional analysis of lncRNA core promoters elucidates rules governing tissue specificity. Genome Res 29: 344–355.

Modzelewski AJ, Gan Chong J, Wang T, He L (2022) Mammalian genome innovation through transposon domestication. Nat Cell Biol 24: 1332–1340.

Necsulea A, Soumillon M, Warnefors M, Liechti A, Daish T, Zeller U, Baker JC, Grützner F, Kaessmann H (2014) The evolution of lncRNA repertoires and expression patterns in tetrapods. Nature 505: 635–640.

Nudelman G, Frasca A, Kent B, Sadler KC, Sealfon SC, Walsh MJ, Zaslavsky E (2018) High resolution annotation of zebrafish transcriptome using long-read sequencing. Genome Res 28: 1415–1425.

O’Neill PK, Forder R, Erill I (2014) Informational requirements for transcriptional regulation. J Comput Biol 21: 373–384.

Patowary A, Purkanti R, Singh M, Chauhan R, Singh AR, Swarnkar M, Singh N, Pandey V, Torroja C, Clark MD et al (2013) A sequence-based variation map of zebrafish. Zebrafish 10: 15–20.

Pauli A, Valen E, Lin MF, Garber M, Vastenhouw NL, Levin JZ, Fan L, Sandelin A, Rinn JL, Regev A et al (2012) Systematic identification of long noncoding RNAs expressed during zebrafish embryogenesis. Genome Res 22: 577–591.

Payne JL, Wagner A (2014) The robustness and evolvability of transcription factor binding sites. Science 343: 875–877.

Pérez-Rico YA, Barillot E, Shkumatava A (2020) Demarcation of Topologically Associating Domains Is Uncoupled from Enriched CTCF Binding in Developing Zebrafish. iScience 23: 101046.

Pillay S, Takahashi H, Carninci P, Kanhere A (2021) Antisense RNAs during early vertebrate development are divided in groups with distinct features. Genome Res 31: 995–1010.

Plaisance I, Chouvardas P, Sun Y, Nemir M, Aghagolzadeh P, Aminfar F, Shen S, Shim WJ, Rochais F, Johnson R et al (2023) A transposable element into the human long noncoding RNA CARMEN is a switch for cardiac precursor cell specification. Cardiovasc Res 119: 1361–1376.

Pollard KS, Hubisz MJ, Rosenbloom KR, Siepel A (2010) Detection of nonneutral substitution rates on mammalian phylogenies. Genome Res 20: 110–121.

Quinlan AR, Hall IM (2010) BEDTools: a flexible suite of utilities for comparing genomic features. Bioinformatics 26: 841–842.

Ranjan G, Sehgal P, Sharma D, Scaria V, Sivasubbu S (2021) Functional long non-coding and circular RNAs in zebrafish. Brief Funct Genomics. doi:10.1093/bfgp/elab014.

Ritter N, Ali T, Kopitchinski N, Schuster P, Beisaw A, Hendrix DA, Schulz MH, Müller-McNicoll M, Dimmeler S, Grote P (2019) The lncRNA Locus Handsdown Regulates Cardiac Gene Programs and Is Essential for Early Mouse Development. Dev Cell 50: 644–657.e8.

Roadmap Epigenomics Consortium, Kundaje A, Meuleman W, Ernst J, Bilenky M, Yen A, Heravi-Moussavi A, Kheradpour P, Zhang Z, Wang J et al (2015) Integrative analysis of 111 reference human epigenomes. Nature 518: 317–330.

Rom A, Melamed L, Gil N, Goldrich MJ, Kadir R, Golan M, Biton I, Perry RB-T, Ulitsky I (2019) Regulation of CHD2 expression by the Chaserr long noncoding RNA gene is essential for viability. Nat Commun 10: 5092.

Santoro F, Mayer D, Klement RM, Warczok KE, Stukalov A, Barlow DP, Pauler FM (2013) Imprinted Igf2r silencing depends on continuous Airn lncRNA expression and is not restricted to a developmental window. Development 140: 1184–1195.

Schertzer MD, Braceros KCA, Starmer J, Cherney RE, Lee DM, Salazar G, Justice M, Bischoff SR, Cowley DO, Ariel P et al (2019) lncRNA-Induced Spread of Polycomb Controlled by Genome Architecture, RNA Abundance, and CpG Island DNA. Mol Cell 75: 523–537.e10.

Sehgal P, Mathew S, Sivadas A, Ray A, Tanwar J, Vishwakarma S, Ranjan G, Shamsudheen KV, Bhoyar RC, Pateria A et al (2021) LncRNA VEAL2 regulates PRKCB2 to modulate endothelial permeability in diabetic retinopathy. EMBO J 40: e107134.

Siepel A, Bejerano G, Pedersen JS, Hinrichs AS, Hou M, Rosenbloom K, Clawson H, Spieth J, Hillier LW, Richards S et al (2005) Evolutionarily conserved elements in vertebrate, insect, worm, and yeast genomes. Genome Res 15: 1034–1050.

Smit AF (1996) The origin of interspersed repeats in the human genome. Curr Opin Genet Dev 6: 743–748.

Smit AF (1999) Interspersed repeats and other mementos of transposable elements in mammalian genomes. Curr Opin Genet Dev 9: 657–663.

Snetkova V, Ypsilanti AR, Akiyama JA, Mannion BJ, Plajzer-Frick I, Novak CS, Harrington AN, Pham QT, Kato M, Zhu Y et al (2021) Ultraconserved enhancer function does not require perfect sequence conservation. Nat Genet 53: 521–528.

Sotero-Caio CG, Platt RN 2nd, Suh A, Ray DA (2017) Evolution and Diversity of Transposable Elements in Vertebrate Genomes. Genome Biol Evol 9: 161–177.

Sutherland GR, Richards RI (1995) Simple tandem DNA repeats and human genetic disease. Proc Natl Acad Sci U S A 92: 3636–3641.

Szcześniak MW, Kubiak MR, Wanowska E, Makałowska I (2021) Comparative genomics in the search for conserved long noncoding RNAs. Essays Biochem 65: 741–749.

Tan H, Onichtchouk D, Winata C (2016) DANIO-CODE: Toward an Encyclopedia of DNA Elements in Zebrafish. Zebrafish 13: 54–60.

Toiber D, Leprivier G, Rotblat B (2017) Long noncoding RNA: noncoding and not coded. Cell Death Discov 3: 16104.

Ulitsky I, Shkumatava A, Jan CH, Sive H, Bartel DP (2011) Conserved function of lincRNAs in vertebrate embryonic development despite rapid sequence evolution. Cell 147: 1537–1550.

Wells JN, Chang N-C, McCormick J, Coleman C, Ramos N, Jin B, Feschotte C (2023) Transposable elements drive the evolution of metazoan zinc finger genes. Genome Res 33: 1325–1339.

Winkler L, Jimenez M, Zimmer JT, Williams A, Simon MD, Dimitrova N (2022) Functional elements of the cis-regulatory lincRNA-p21. Cell Rep 39: 110687.

Yang H, Luan Y, Liu T, Lee HJ, Fang L, Wang Y, Wang X, Zhang B, Jin Q, Ang KC et al (2020) A map of cis-regulatory elements and 3D genome structures in zebrafish. Nature 588: 337–343.

Yap KL, Li S, Muñoz-Cabello AM, Raguz S, Zeng L, Mujtaba S, Gil J, Walsh MJ, Zhou M-M (2010) Molecular interplay of the noncoding RNA ANRIL and methylated histone H3 lysine 27 by polycomb CBX7 in transcriptional silencing of INK4a. Mol Cell 38: 662–674.

Yin Y, Yan P, Lu J, Song G, Zhu Y, Li Z, Zhao Y, Shen B, Huang X, Zhu H et al (2015) Opposing Roles for the lncRNA Haunt and Its Genomic Locus in Regulating HOXA Gene Activation during Embryonic Stem Cell Differentiation. Cell Stem Cell 16: 504–516.

Yuan J, Zhang X, Wang M, Sun Y, Liu C, Li S, Yu Y, Gao Y, Liu F, Zhang X et al (2021) Simple sequence repeats drive genome plasticity and promote adaptive evolution in penaeid shrimp. Commun Biol 4: 186.

Yuan W, Jiang S, Sun D, Wu Z, Wei C, Dai C, Jiang L, Peng S (2019) Transcriptome profiling analysis of sex-based differentially expressed mRNAs and lncRNAs in the brains of mature zebrafish (Danio rerio). BMC Genomics 20: 830.

Zhou Q, Jiang Y, Cai C, Li W, Leow MK-S, Yang Y, Liu J, Xu D, Sun L (2023) Multidimensional conservation analysis decodes the expression of conserved long noncoding RNAs. Life Sci Alliance 6. doi:10.26508/lsa.202302002.

