## Supplementary material for "Syntenic lncRNAs exhibit DNA regulatory functions with sequence evolution": Supply Fig

### Supplementary files.

Supplementary Table 1:-

| Tissue | Timepoint | File |
|---|---|---|
| stomach | 0 | ENCSR178GUS |
| forebrain | 0 | ENCSR362AIZ |
| kidney | 0 | ENCSR173PJN |
| liver | 0 | ENCSR096STK |
| lung | 0 | ENCSR982MRY |
| midbrain | 0 | ENCSR719NAJ |
| heart | 0 | ENCSR526SEX |
| intestine | 0 | ENCSR331XCE |
| hindbrain | 0 | ENCFF775PFB |
| hindbrain | 0 | ENCFF263FJA |
| forebrain | 11.5 | ENCSR160IIN |
| limb | 11.5 | ENCSR541XZK |
| heart | 11.5 | ENCSR691OPQ |
| liver | 11.5 | ENCSR284AMY |
| neural_tube | 11.5 | ENCSR337FYI |
| midbrain | 11.5 | ENCSR307BCA |
| hindbrain | 11.5 | ENCFF400JIC |
| hindbrain | 11.5 | ENCFF658AUO |
| forebrain | 12.5 | ENCSR647QBV |
| limb | 12.5 | ENCSR750YSX |
| liver | 12.5 | ENCSR648YEP |
| hindbrain | 12.5 | ENCSR420QTO |

| Tissue | Timepoint | File |
|---|---|---|
| stomach | 0 | ENCSR178GUS |
| forebrain | 0 | ENCSR362AIZ |
| kidney | 0 | ENCSR173PJN |
| liver | 0 | ENCSR096STK |
| lung | 0 | ENCSR982MRY |
| midbrain | 0 | ENCSR719NAJ |
| heart | 0 | ENCSR526SEX |
| intestine | 0 | ENCSR331XCE |
| midbrain | 12.5 | ENCSR908JWT |
| heart | 12.5 | ENCSR150CUE |
| neural_tube | 12.5 | ENCFF494DTT |
| neural_tube | 12.5 | ENCFF964TAM |
| hindbrain | 13.5 | ENCSR921PRX |
| midbrain | 13.5 | ENCSR792RJV |
| forebrain | 13.5 | ENCSR970EWM |
| neural_tube | 13.5 | ENCFF789LMJ |
| neural_tube | 13.5 | ENCFF028DKY |
| heart | 13.5 | ENCFF202UVT |
| heart | 13.5 | ENCFF245ZGS |
| limb | 13.5 | ENCFF703ILV |
| limb | 13.5 | ENCFF466OJM |
| liver | 13.5 | ENCFF661DXP |
| liver | 13.5 | ENCFF938KFW |
| lung | 14.5 | ENCSR039ADS |

| **Tissue** | **Timepoint** | **File** |
|---|---|---|
| stomach | 0 | ENCSR178GUS |
| forebrain | 0 | ENCSR362AIZ |
| kidney | 0 | ENCSR173PJN |
| liver | 0 | ENCSR096STK |
| lung | 0 | ENCSR982MRY |
| midbrain | 0 | ENCSR719NAJ |
| heart | 0 | ENCSR526SEX |
| intestine | 0 | ENCSR331XCE |
| midbrain | 14.5 | ENCSR343YLB |
| kidney | 14.5 | ENCSR504GEG |
| stomach | 14.5 | ENCSR290RRR |
| limb | 14.5 | ENCSR216NEG |
| neural_tube | 14.5 | ENCSR928OXI |
| forebrain | 14.5 | ENCSR185LWM |
| intestine | 14.5 | ENCSR932TRU |
| hindbrain | 14.5 | ENCSR559TRB |
| liver | 14.5 | ENCSR867YNV |
| heart | 14.5 | ENCFF196SUL |
| heart | 14.5 | ENCFF080KZO |
| limb | 15.5 | ENCSR830IVQ |
| lung | 15.5 | ENCSR457RRW |
| midbrain | 15.5 | ENCSR557RMA |
| forebrain | 15.5 | ENCSR752RGN |
| intestine | 15.5 | ENCSR370SFB |

| **Tissue** | **Timepoint** | **File** |
|---|---|---|
| stomach | 0 | ENCSR178GUS |
| forebrain | 0 | ENCSR362AIZ |
| kidney | 0 | ENCSR173PJN |
| liver | 0 | ENCSR096STK |
| lung | 0 | ENCSR982MRY |
| midbrain | 0 | ENCSR719NAJ |
| heart | 0 | ENCSR526SEX |
| intestine | 0 | ENCSR331XCE |
| kidney | 15.5 | ENCSR062VTB |
| liver | 15.5 | ENCSR611PTP |
| neural_tube | 15.5 | ENCFF346PRA |
| neural_tube | 15.5 | ENCFF301NKF |
| heart | 15.5 | ENCFF027WCY |
| heart | 15.5 | ENCFF836AFL |
| stomach | 15.5 | ENCFF376OIM |
| stomach | 15.5 | ENCFF537GAJ |
| hindbrain | 15.5 | ENCFF675CWB |
| hindbrain | 15.5 | ENCFF894AVY |
| stomach | 16.5 | ENCSR466KZY |
| lung | 16.5 | ENCSR992WBR |
| hindbrain | 16.5 | ENCSR285WZV |
| forebrain | 16.5 | ENCSR080EVZ |
| liver | 16.5 | ENCSR826HIQ |
| heart | 16.5 | ENCSR020DGG |

| Tissue | Timepoint | File |
|---|---|---|
| stomach | 0 | ENCSR178GUS |
| forebrain | 0 | ENCSR362AIZ |
| kidney | 0 | ENCSR173PJN |
| liver | 0 | ENCSR096STK |
| lung | 0 | ENCSR982MRY |
| midbrain | 0 | ENCSR719NAJ |
| heart | 0 | ENCSR526SEX |
| intestine | 0 | ENCSR331XCE |
| kidney | 16.5 | ENCSR537GNQ |
| midbrain | 16.5 | ENCSR367ZPZ |
| midbrain | 16.5 | ENCFF861MLR |
| intestine | 16.5 | ENCFF410BTA |
| intestine | 16.5 | ENCFF411WZW |

**Supplementary Figures:-**

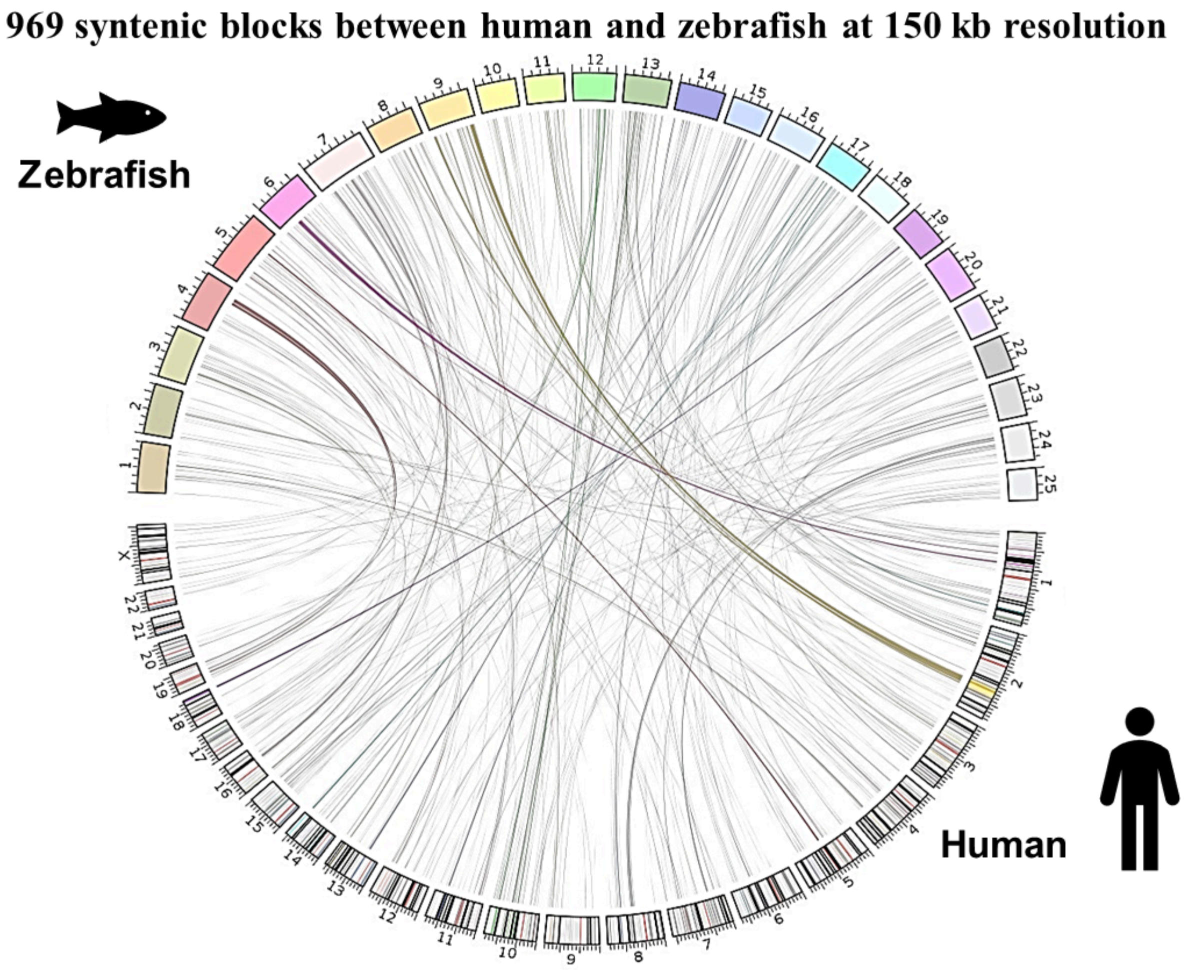


**Supplementary Figure 1 :-** Circos plot representing 969 conserved syntenic blocks between human and zebrafish chromosomes.

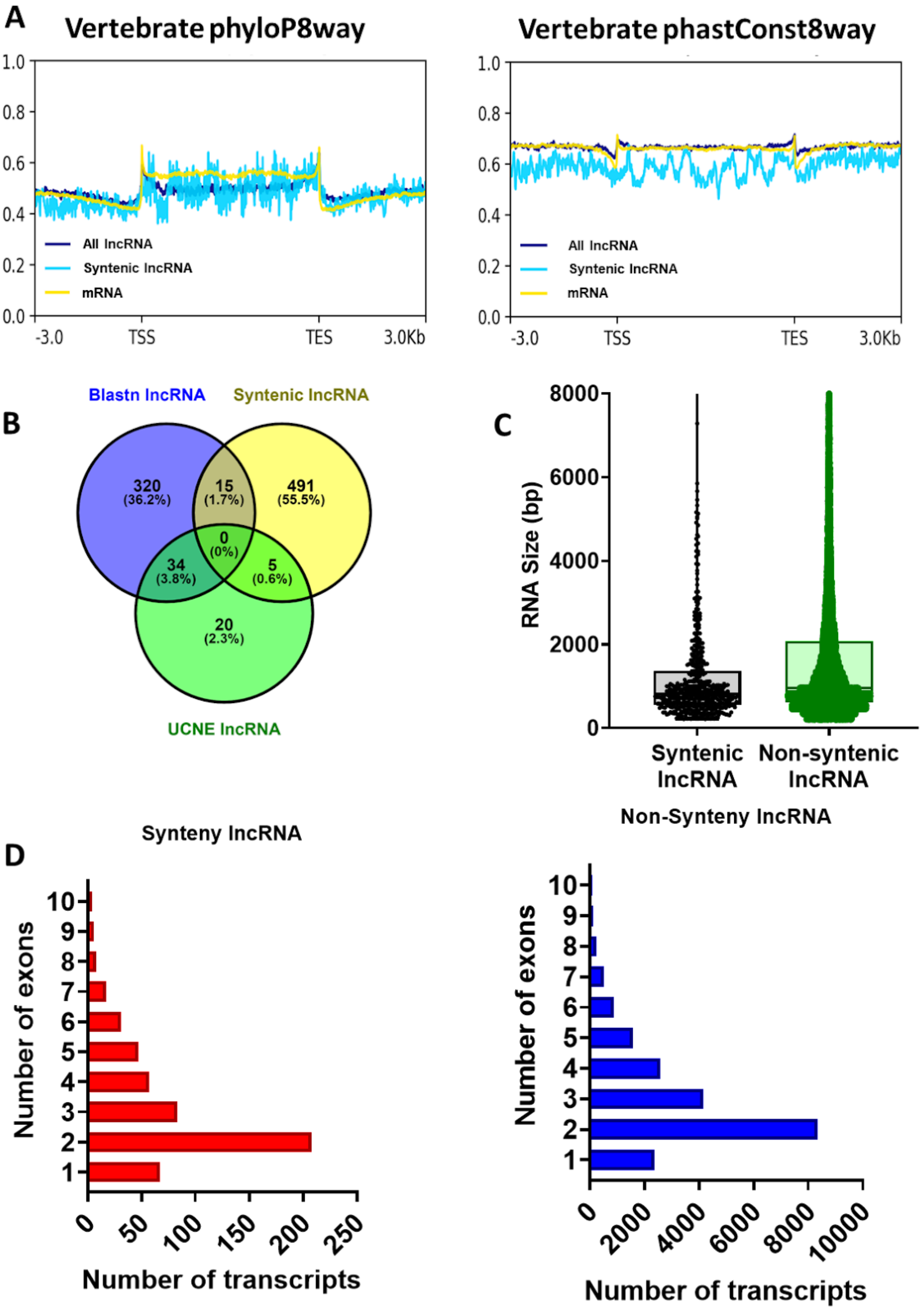


**Supplementary Figure 2:** [A] Profile plot of PhyloP8way and PhastCon8 way for all the lncRNAs, syntenic lncRNAs and mRNAs of zebrafish.[B] Venn plot of all the lncRNAs showing overlap in type of conservation observed between human and zebrafish. [C] Box plot representing the length of the syntenic lncRNAs and non-syntenic lncRNAs. [D] Bar plot representing the number of exon present in the syntenic lncRNA and non syntenic lncRNA.

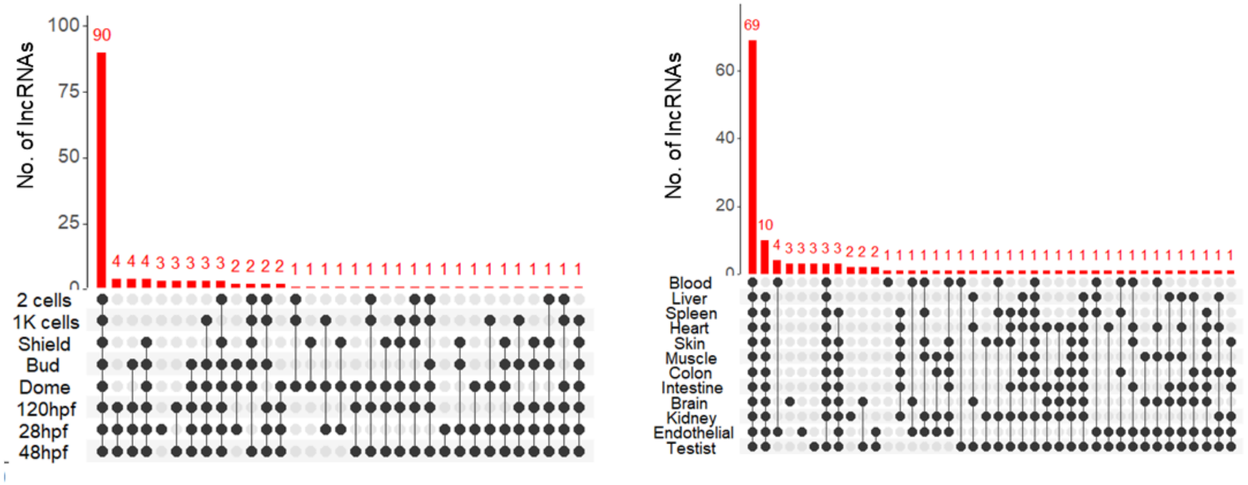


**Supplementary Figure 3:-** Upset plot representing the overlap of lncRNAs expression across different developmental stages and in different adult tissues of zebrafish.

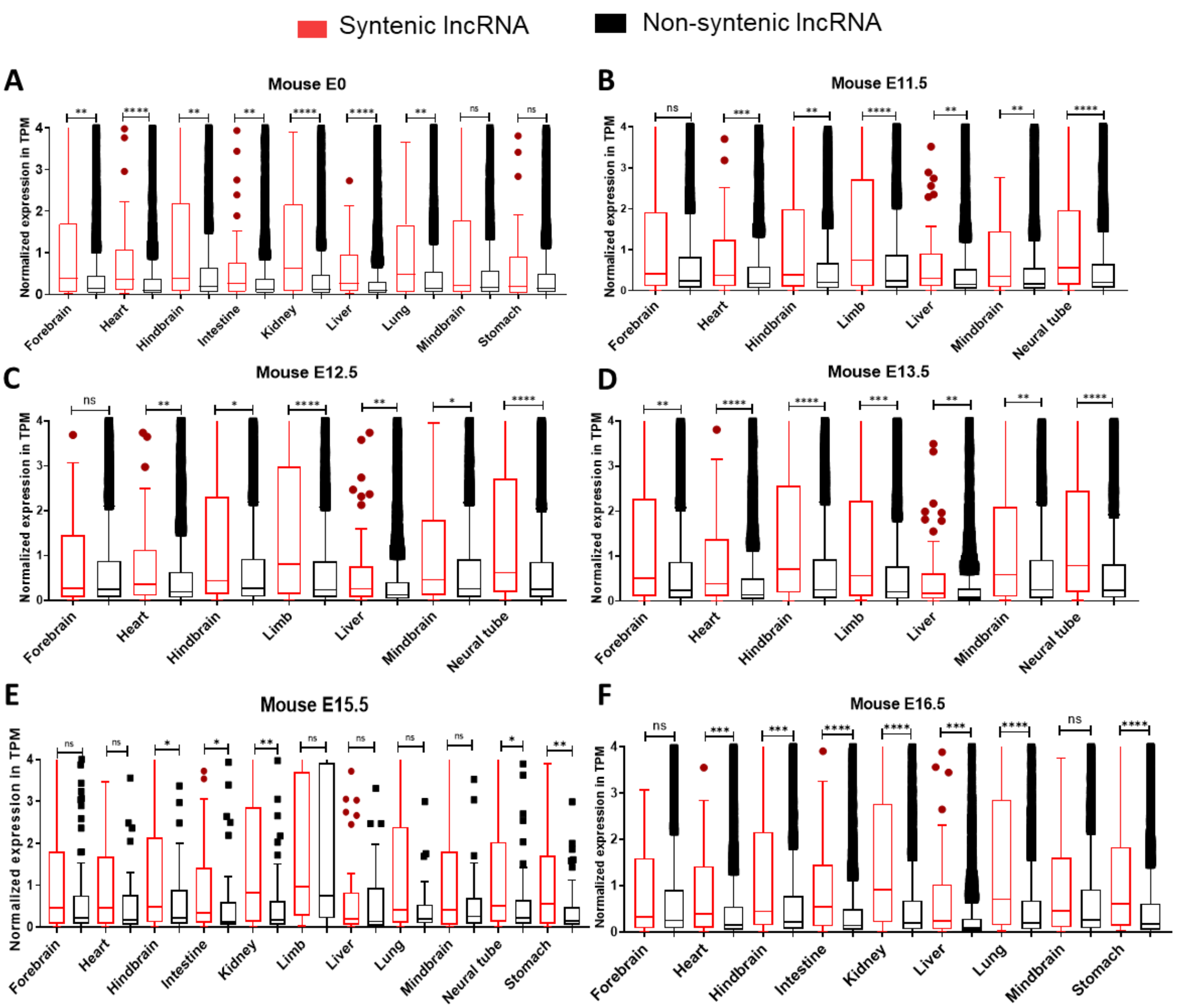


**Supplementary Figure 4:-**Expression profile of syntenic and non-syntenic lncRNAs across early development stage of mouse [A]E0; [B] E11.5 ; [C]E12.5; [D]E13.5; [E]E15.5; and [F] E16.5 stage. ns- not significant; **** P<0.0001; *** P<0.001; ** P<0.01;* P<0.05 (Mann-Whitney U).

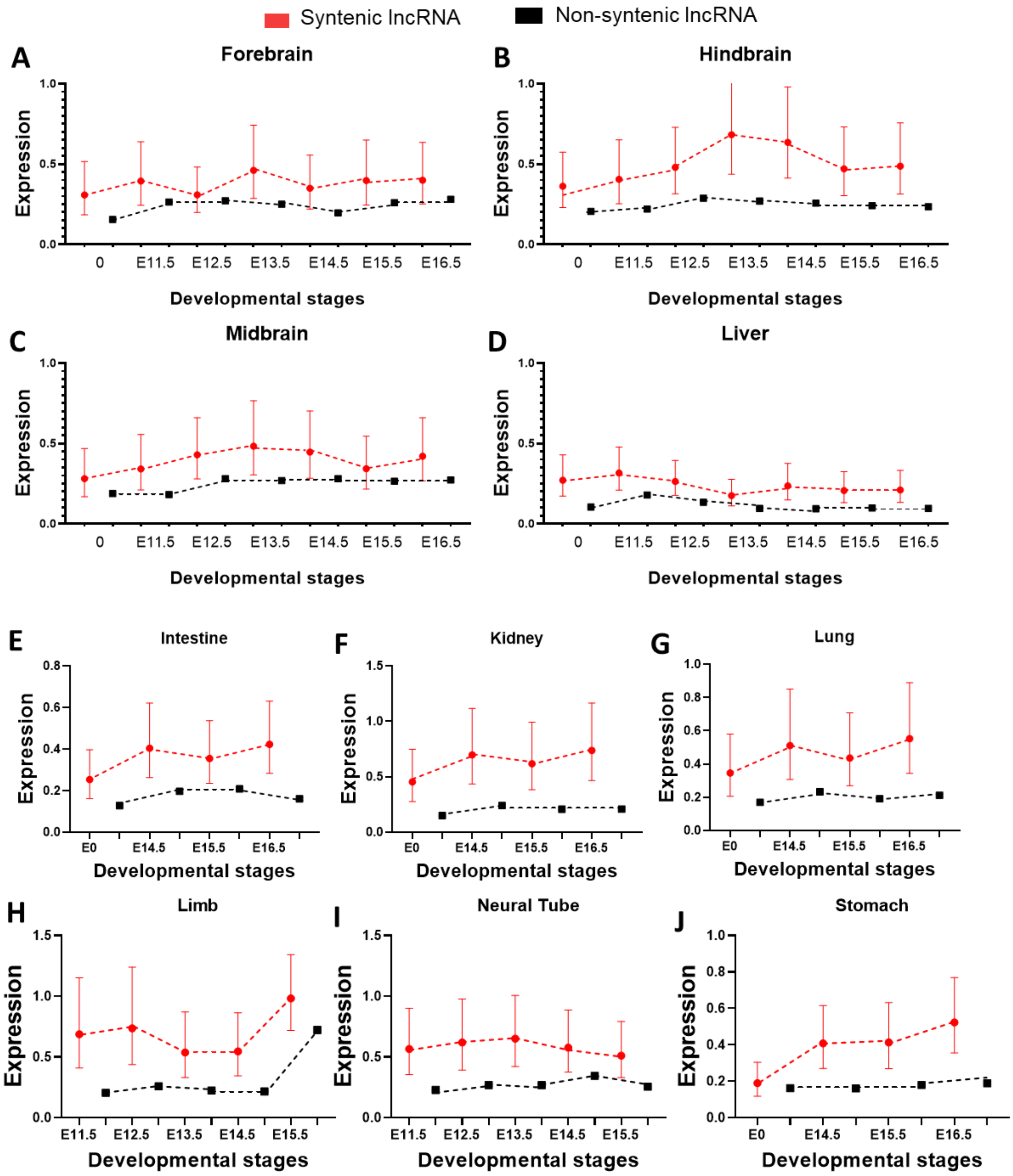


**Supplementary Figure 5:-**Dynamic expression profile of mouse tissues represented as geometric mean with 95% CI for syntenic and non-syntenic lncRNAs.[A]Forebrain; [B]Hindbrain; [C]Midbrain; [D]Liver; [E]Intestine; [F]Kidney; [G]Lung; [H]Limb; [I]Neural Tube; and [J]Stomach.

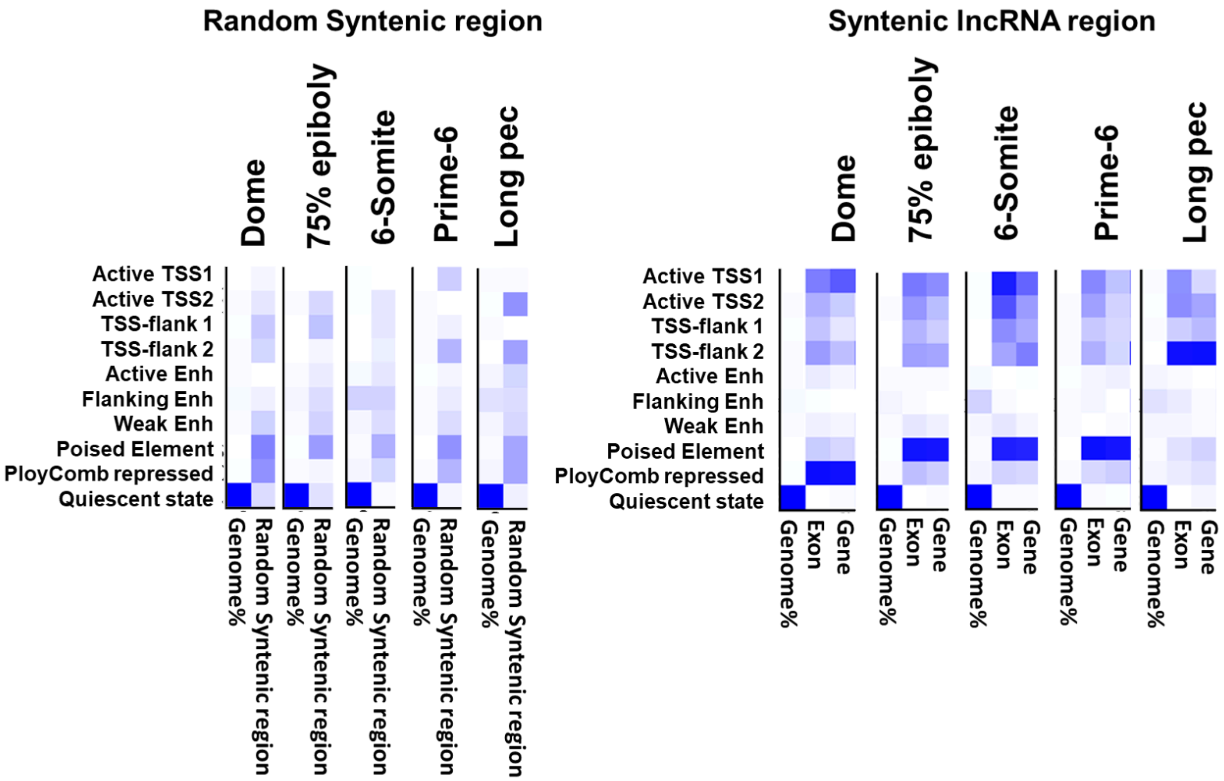


**Supplementary Figure 6:** Overlap enrichment of syntenic and random size matched syntenic region across chromatin states of 5 zebrafish developmental stages. The Chromatin state is represented on the y-axis, and the region is classified on the x-axis. Color represents the enrichment of the chromatin state. Highlighted with red represented chromatin state in focus. The heatmap has a specific color scale wherein white will correspond to 0 and the darkest color will be the maximum value

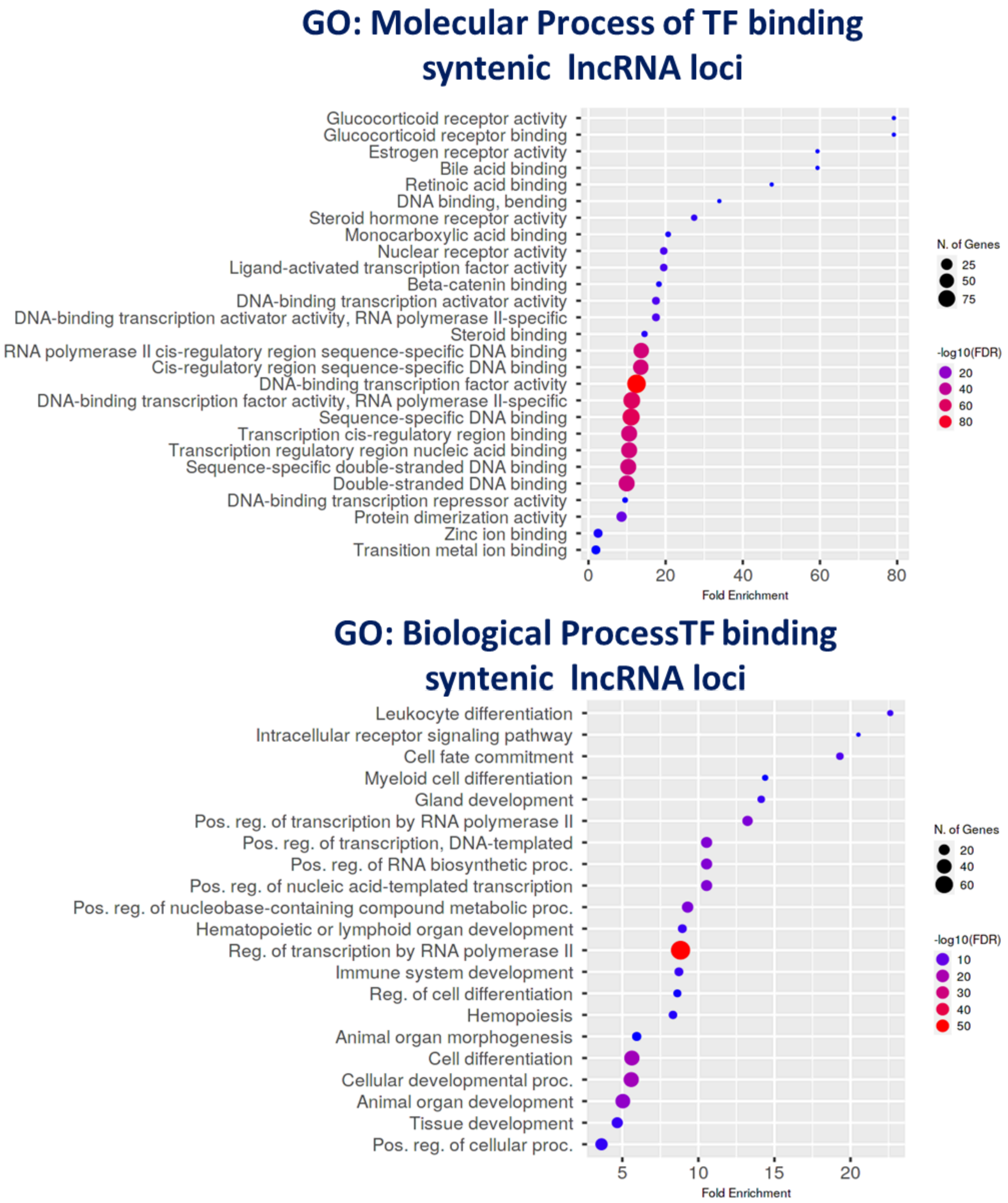


**Supplementary Figure 7 :** Gene ontology of molecular and biological process for identified significant transcription factor binding on the syntenic lncRNA locus.

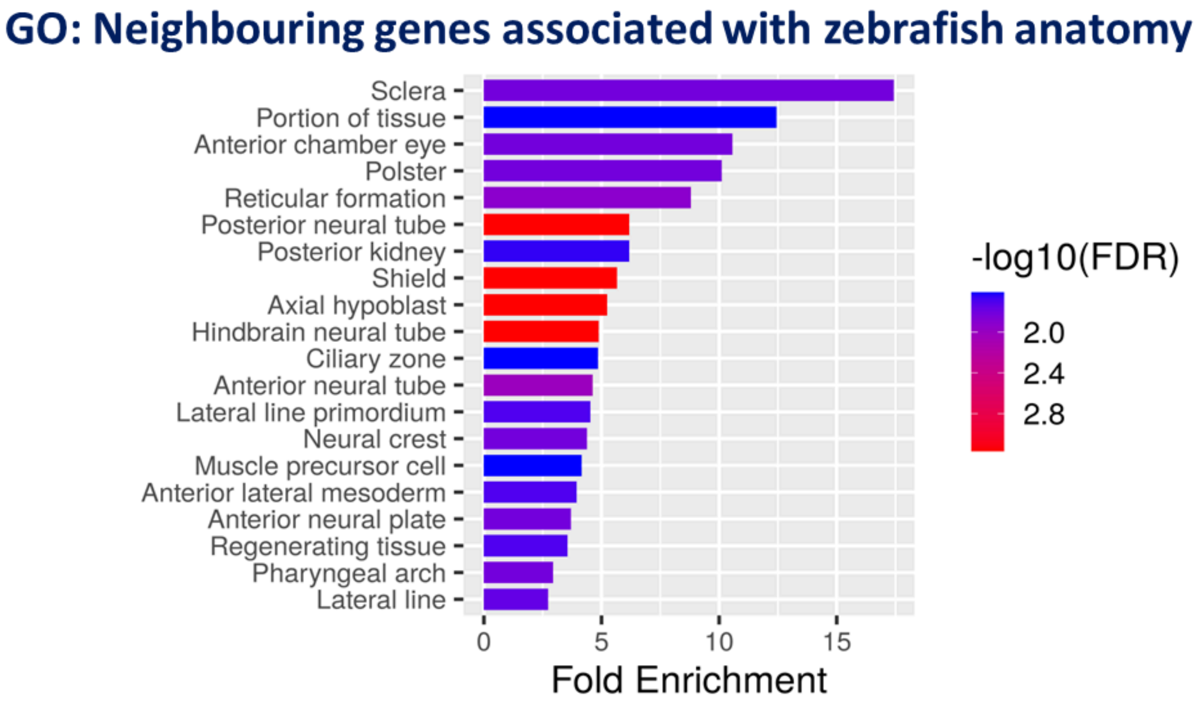


**Supplementary Figure 8 :-** Gene ontology for the syntenic lncRNAs neighboring protein-coding gene associated with zebrafish anatomy

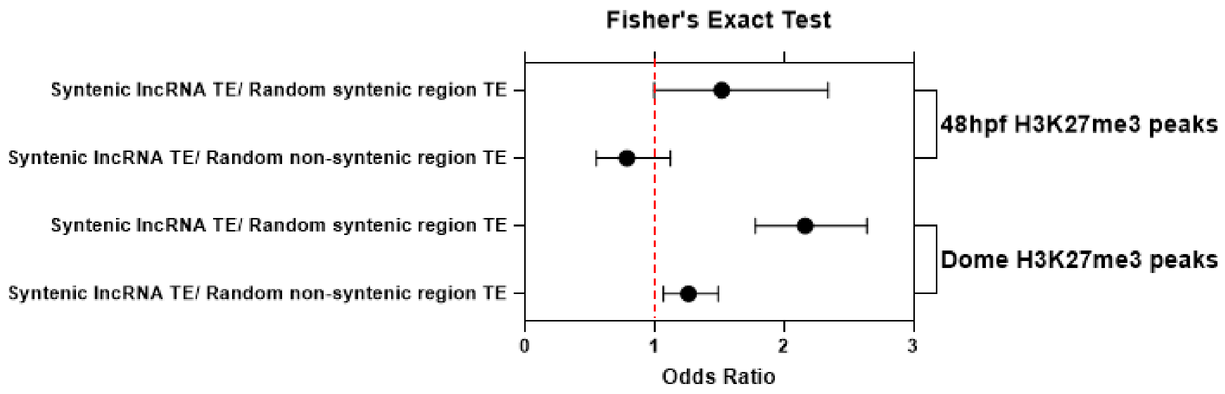


**Supplementary Figure 9:** Forest plot showing the odds ratios and 95% confidence intervals for overlap between lncRNAs, TEs, and H3K27me3 peaks in syntenic lncRNAs, Random Syntenic regions, and non-syntenic lncRNAs in Dome and 48 hpf zebrafish.(Fisher's t-Test)

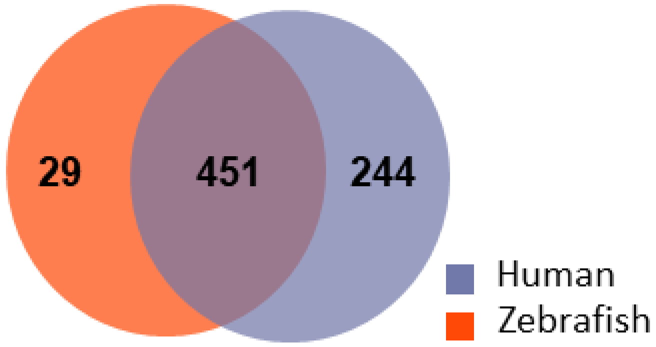


**Supplementary Figure 10 :-** Venn plot representing unique and overlap transcription factors binding on the *ZFLNCG00003* lncRNA zebrafish and *ENSG00000283828* lncRNA in human from JASPER 2022 database.
